# Harmonized Protocol for Segmentation of the Hippocampal Tail on High-Resolution *in vivo* MRI from the Hippocampal Subfields Group (HSG)

**DOI:** 10.1101/2025.11.20.689476

**Authors:** Robin de Flores, Kelsey L Canada, Thackery Brown, Nicole J Gervais, Anne Maass, Gustaf Rådman, Jonathan Shine, Hannah L. Tucker, Eóin N. Molloy, Jenna N. Adams, Michelle Baca Reinke, Arnold Bakker, David Berron, Marshall A Dalton, Kristen M Kennedy, Renaud LaJoie, Susanne G. Mueller, Noa Ofen, Rosanna K Olsen, Naftali Raz, Tracy Riggins, Karen M Rodrigue, Craig Stark, Lei Wang, Laura E.M. Wisse, Paul A. Yushkevich, Valerie A. Carr, Ana M. Daugherty, the Alzheimer’s Disease Neuroimaging Initiative, the Hippocampal Subfields Group

**Affiliations:** Normandie Univ, UNICAEN, INSERM, UA20 Neuropresage, 14000 Caen, France; University of Massachusetts Amherst; School of Psychology, Center for Research and Education in Navigation (CRaNE), Georgia Institute of Technology; University of Groningen; German Center for Neurodegenerative Diseases (DZNE), Magdeburg; Lund University; German Center for Neurodegenerative Diseases (DZNE), Magdeburg, Germany; Department of Clinical Sciences, Lund University; Department of Neurobiology and Behavior, University of California Irvine; San Jose State University; Department of Psychiatry and Behavioral Sciences, Johns Hopkins University, Baltimore, MD, USA; University of Sydney; Center for Vital Longevity, School of Behavioral and Brain Sciences, The University of Texas at Dallas; Memory and Aging Center, UCSF Weill Institute for Neurosciences, Department of Neurology, University of California, San Francisco, California; Department of Radiology and Biomedical Imaging, University of California, San Francisco; Center for Vital Longevity, School of Behavioral and Brain Sciences, University of Texas at Dallas; Baycrest Academy for Research and Education, University of Toronto; Stony Brook University, Max Planck Institute for Human Development, Berlin, Germany; University of Maryland; University of California Irvine; Rotman Research Institute, Baycrest Academy for Research and Education, University of Toronto; Department of Clinical Sciences Lund, Lund University; Department of Radiology, University of Pennsylvania Perelman School of Medicine; Wayne State University

**Author notes:** V. A. Carr and A. M. Daugherty should be considered joint senior authors. Data used in preparation of this article were obtained from the Alzheimer’s Disease Neuroimaging Initiative (ADNI) database (adni.loni.usc.edu). As such, the investigators within the ADNI contributed to the design and implementation of ADNI and/or provided data but did not participate in analysis or writing of this report. A complete listing of ADNI investigators can be found at:http://adni.loni.usc.edu/wp-content/uploads/how_to_apply/ADNI_Acknowledgement_List.pdf.

## Abstract

The hippocampus is a heterogeneous structure with cytoarchitectonically distinct subfields that exhibit heterogeneous lifespan trajectories and are differentially susceptible to diseases. Advances in high-resolution imaging have accelerated research on these structures, yet variability in segmentation protocols limits cross-study comparability. The Hippocampal Subfields Group (HSG) is an international consortium addressing this challenge by developing a reliable, accessible, and freely available segmentation protocol for high-resolution T2-weighted 3 tesla MRI scans (http://www.hippocampalsubfields.com). Here, we present the harmonized protocol for the posterior portion of the hippocampus (the “tail”), complementing the previously established “body” protocol, and with an anterior “head” protocol under development. The tail protocol provides standardized definitions of the external boundaries for the posterior-most extent of the hippocampus, facilitating consistent segmentation from surrounding tissues. The research community was extensively involved through an online survey that incorporated comprehensive protocol details, feasibility assessments, tutorial videos, and illustrative segmentations. Through this collaborative process, consensus emerged to exclude subfield labeling in the hippocampal tail due to limited visibility of internal landmarks and substantial anatomical variability in this region. All proposed boundary guidelines were deemed clear and agreed upon via a Delphi procedure. The harmonized tail protocol has high intra-(Averaged ICC(2,1) > 0.98; Averaged Dice Similarity Coefficient = 0.92) and inter-rater reliability (Averaged ICC(2,k) > 0.98; Averaged Dice Similarity Coefficient = 0.86) and offers a practical framework for replicable segmentation. By establishing standardized guidelines, this protocol enhances comparability of findings across developmental, aging, and clinical research and is compatible with ongoing technological advances.

## 1. Introduction

The hippocampal formation is a neural nexus with a role in a wide range of important cognitive functions, such as spatial processing (Moser et al., 2008; Fenton, 2024), episodic memory (Carr et al., 2010; Moscovitch et al., 2016; Kolibius et al., 2025), and imagination (Buckner, 2010). However, this important brain region is also particularly susceptible to a range of neuropathological influences that feature prominently in neurological and psychiatric diseases including Alzheimer’s disease (AD), schizophrenia, and depression (Geuze et al., 2005; Small et al., 2011; Bartsch and Wulff, 2015). This confluence of importance and vulnerability makes the hippocampus a hotspot for neuroimaging research. Importantly, the hippocampus is a heterogeneous structure, comprising cytoarchitectonically distinct subfields that differ in function, connectivity with other brain areas and susceptibility to diseases (Small et al., 2011; Aggleton, 2012; Duvernoy et al., 2013; Maruszak and Thuret, 2014; Wisse et al., 2025). Neuroanatomical consensus identifies five subfields: the subicular complex (Sub), dentate gyrus (DG), and Cornu Ammonis (CA) sectors 1 through 3. Some neuroanatomists add a CA4 region (Ding, 2013; Duvernoy et al., 2013; Palomero-Gallagher et al., 2020), while others refer to this region as hilus (West and Gundersen, 1990) or part of CA3 (Insausti and Amaral, 2012). Additionally, the hippocampus can be also divided along an antero-posterior axis, analogous to the ventral-dorsal gradient in rodents, and is generally considered a tripartite structure, including the hippocampal head, body, and tail (Aggleton, 2012; Poppenk et al., 2013; Strange et al., 2014; Plachti et al., 2019; Genon et al., 2021).

In the 2000s, technical and methodological advances enabled the development of high-resolution T2-weighted *in vivo* imaging, which has been increasingly used to assess age- or disease-related variations in hippocampal subfields (Mueller et al., 2007; de Flores et al., 2015; Grande et al., 2023; Sun et al., 2023; Yushkevich et al., 2024; Zilioli et al., 2025). The rapid accumulation of empirical studies in this field was accompanied by the development of numerous protocols for *in vivo* segmentation and labeling of human hippocampal subfields on MRI. However, variations in anatomical nomenclature and segmentation heuristics have created challenges for synthesizing and interpreting findings across studies, thereby hindering scientific progress and clinical application (Yushkevich et al., 2015).

To address these challenges, the Hippocampal Subfields Group (HSG - https://hippocampalsubfields.com/) was established to develop a harmonized protocol for segmenting hippocampal subfields and the medial temporal lobe (MTL) cortex. This protocol is designed to be applicable across the lifespan and in populations with diverse diseases. The HSG currently includes more than 250 scientists from over 30 countries, spanning all career stages and levels of experience. The group has produced several collaborative publications, including a comparison of 21 protocols to assess labeling disagreements (Yushkevich et al., 2015), an overview of the HSG’s goals and structure (Wisse et al., 2017), a progress update on protocol development for the hippocampal body (Olsen et al., 2019), a guide to quality control for high-resolution T2-weighted MRI (Canada et al., 2024), a comparison of histological delineations of MTL cortices by four independent neuroanatomy laboratories (Wuestefeld et al., 2024) and, most recently, the harmonized protocol for segmenting subfields in the hippocampal body in high-resolution T2-weighted MRI (Daugherty et al., 2025). The HSG opted to accelerate progress by developing separate protocols for the hippocampal head, body and tail in parallel across different working groups.

The hippocampal tail (HT), and more broadly the posterior portion of the hippocampus, has been associated with episodic memory, particularly retrieval processes (Lepage et al., 1998; Kim, 2015), and with spatial navigation (Strange et al., 2014; Brunec et al., 2019). Furthermore, the posterior hippocampus may be more sensitive to aging and age-associated cognitive decline than the anterior region (Kalpouzos et al., 2009; Chauveau et al., 2021), although there is no consensus on this matter (Chen et al., 2010; Ta et al., 2012). These functional and structural characteristics, and the discrepant results across studies underscore the importance of developing a consistent and valid protocol for HT segmentation.

In this context, the present study outlines the procedures used to develop the harmonized protocol for labeling the HT, along with the supportive validation evidence. We comprehensively describe the harmonized HT protocol which was developed and validated using high-resolution T2-weighted MRI (0.4 × 0.4 mm² in-plane) that enables visualization of the anatomical details necessary for accurate segmentation (Wisse et al., 2017; Canada et al., 2024). The HT protocol presented here, along with the recently published body protocol (Daugherty et al., 2025) and the upcoming head protocol, will form the Harmonized Hippocampal Subfields Segmentation Protocol.

### Overview of Protocol Development and Validation Process

Like the body protocol, the HT protocol was developed using a Delphi procedure inspired by the EADC (European Alzheimer’s Disease Consortium)-ADNI (Alzheimer’s Disease Neuroimaging Initiative) Harmonized Protocol (HarP) (Boccardi et al., 2015). This iterative process incorporates expert input from the research community, helping to integrate diverse approaches into a comprehensive, cohesive protocol. It also fosters community support and encourages its implementation.

Unlike the hippocampal body and head, the HT is not segmented into its constituent subfields. The application of a single label for HT was motivated by two considerations. First, it is much more difficult to visualize the inner structure of the HT on MR images, making subfield segmentation based on visible landmarks such as the Stratum Radiatum, Lacunosum, and Moleculare (SRLM) less feasible (Figure 1). This is due to partial volume effects, most likely created by the region wrapping through the acquisition plane, that result in greater blurring of the features in the tail than in the head and body.

**Figure 1:**
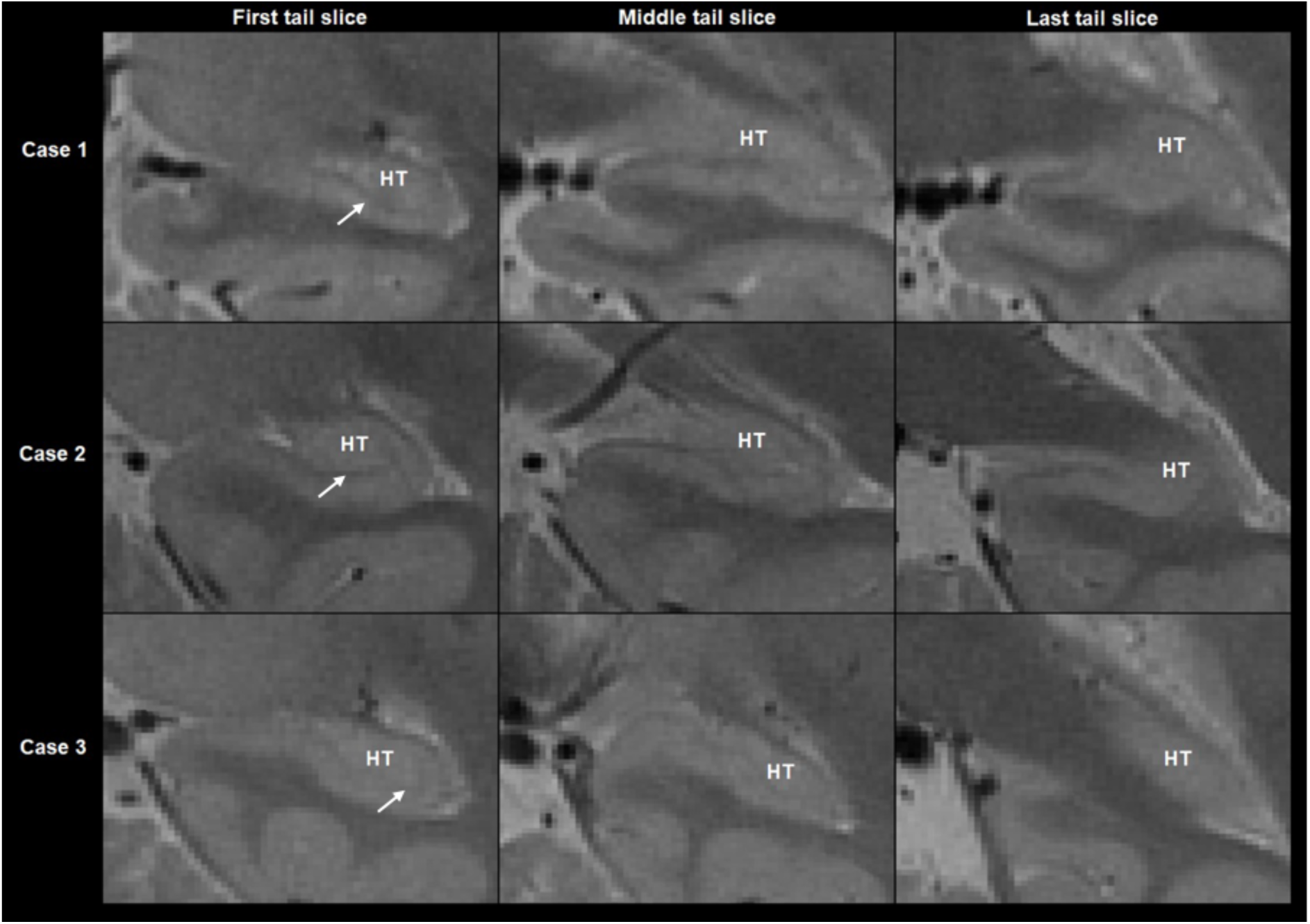
Visualization of tail slices on T2-weighted coronal sections across three cases illustrating the lack of visible inner structures across the HT such as the SRLM (dark band, see arrow).

Second, defining subfields in the HT is further hampered by substantial individual differences in the appearance of the tail on coronal sections due to variations in its curvature (Adler et al., 2018). Consequently, the appearance and location of the subfields in the tail is highly variable in the coronal plane (Figure 2), and it is difficult to specify a single protocol that would fit all variations. Reslicing MR images along the curvature of the tail with varying orientation relative to the long axis of the hippocampus can yield the level of detail in shape of the structure that is comparable to those of the body (de Flores et al., 2020). However, the anisotropic resolution in the z-plane of typical T2-images acquired with 1-3 mm slice thickness makes reslicing unfeasible.

**Figure 2:**
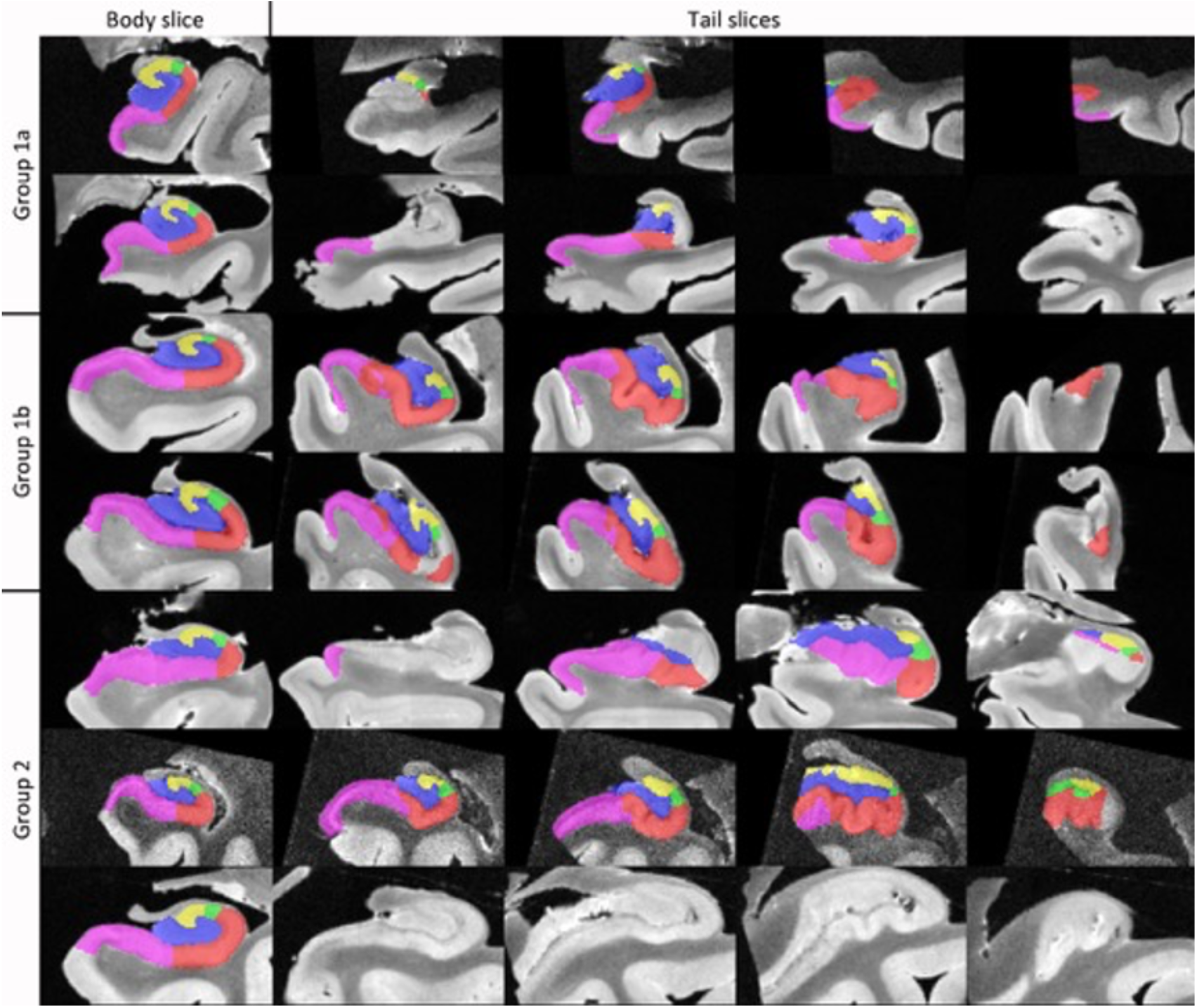
Figure adopted from (de Flores et al., 2020) showing comparisons of a body slice to several tail slices, highlighting the varied curvature and subfield appearance in the tail across different specimens. Group 1a: a body-like shape can be observed in the hippocampal tail. Group 1b: a body-like shape where the tail curves in a superior direction. Group 2: The shape of the tail is not body-like and the tail curves in a medial direction. In the last case no histology-based segmentations were available in the posterior portion of the hippocampus, however, a similar shape can be observed based on the raw MR image. Colors: yellow: CA3, green: CA2, red: CA1, pink: subiculum, blue: DG.

Therefore, the HT protocol does not rely on the histological dataset used by the HSG in developing other protocols (Olsen et al., 2019; Daugherty et al., 2025). Otherwise, the development of this protocol follows the same general procedure as the other HSG protocols: Landmarks and guidelines for delineating the outer boundaries of the hippocampal tail were identified in MRI space, while considering anatomical atlases (Mai et al., 2008; Duvernoy et al., 2013), followed by an initial feasibility check. From this point, feedback was collected using the Delphi procedure, the response of which was used to repeat the protocol development steps, until reaching consensus and performing a formal reliability analysis (Wisse et al., 2017; Olsen et al., 2019; Daugherty et al., 2025).

## 2. Methods

### 2.1. Rule development for ranging and outer boundaries on MRI

We developed a step-by-step protocol for identifying anatomical landmarks and defining HT boundaries. The process began with identifying landmarks on *in vivo* structural MRI that mark the posterior limit of the hippocampal tail. Note that the anterior limit of the HT, which coincides with the posterior limit of the hippocampal body, was defined previously (Olsen et al., 2019). Next, we established the outer boundaries of the HT to clearly separate hippocampal tissue from surrounding structures and CSF. This protocol was created by the four members of the HSG Tail Working Group (RdF, NG, AM and JS) and was initiated during an open meeting held in 2018 in Magdeburg, Germany, with additional input from attending members of the broader HSG community.

### 2.2. Initial check for feasible reliability

To prepare for the Delphi procedure, we assessed the feasibility of the HT protocol using a small, representative image set to obtain reliability estimates and gather qualitative feedback from raters. Two expert raters with at least 4 years of experience segmenting the hippocampus participated in the feasibility assessment. One was part of the HT working group and the other was completely naive to the protocol prior to training. Training included detailed documentation with example image tracings, a 1-hour introductory training session (via Zoom), followed by prescribed practice and then an additional 1-hour of individualized feedback (via Zoom).

The dataset for the feasibility testing included *in vivo* MRI brain scans of typically developing children (n = 2, both male, age 9 and 15 years), healthy adults (n = 1, male age 66), and a patient with dementia of the Alzheimer’s type collected by ADNI (n = 1, female age 70). A self-report questionnaire was distributed to compile information regarding the rater’s experience, i.e. their understanding and agreement with the proposed protocol. The raters responded using Likert scales as well as open response to gather qualitative information regarding the protocol feasibility. All segmentations were produced using ITK-SNAP (Yushkevich et al., 2006), as this is a free, easily accessible and open-sourced tool. This harmonized protocol was, however, designed to be compatible with any manual segmentation software.

### 2.3. Feedback collection via Delphi procedure

As with development of the body protocol (Daugherty et al., 2025), a Delphi-panel consisting of experts was provided with quantitative and qualitative evidence for the landmarks and boundary definitions to be used for segmentation of the HT. Panelists were recruited via an open call to the HSG member email-list, website, and over various social media platforms. The questionnaire was completed collaboratively by each participating lab, and these responses were subsequently anonymized. When completing the questionnaire, panelists first answered a series of questions pertaining to the populations they typically work with, the types of scan sequences they typically use, and their degree of experience with segmenting the hippocampal tail. Then, they evaluated each rule using a 9-point Likert scale to assess both agreement with the rule (1 = do not agree at all; 9 = fully agree) and clarity of the rule (1 = extremely unclear, requiring major revisions; 9 = extremely clear, requiring no changes) in an iterative and recursive process until consensus was reached for the full segmentation protocol. Panelists were able to opt out of voting on a particular rule if they had no opinion. Consensus on a given rule was declared when the number of agreement responses (Likert rating 6-9) was statistically greater than the number of disagreement responses (Likert rating 1-5) (Daugherty et al., 2025).

The Delphi-panel responses were obtained through Qualtrics (https://www.qualtrics.com; Qualtrics, Provo, UT) and collected between December 2023 and March 2024. A compressed file with supplemental material was provided as an accompaniment to the Qualtrics questionnaire. This contained example HT segmentations, the MRI scans included in the feasibility testing, a spreadsheet with feasibility data, a video file with demonstration of the protocol and a 31-page supplemental document. Delphi panelists were asked to review and interact with the material, encouraged to apply the protocol to segment one or more of the sample MRI scans available, before completing the questionnaire. Completion of the full questionnaire was estimated to take around 30 minutes, excluding preparation time with the material at hand and segmentation practice.

### 2.4. Formal reliability analysis of the consensus protocol

Following consensus through the Delphi procedure, three raters conducted a formal reliability analysis of the final segmentation protocol. The raters were the same individuals who participated in the reliability testing of the hippocampal body protocol. Their experience with hippocampal segmentation varied: two were experts with over 4 years of manual segmentation experience, while one had no prior experience with segmenting the hippocampal tail. Their familiarity with the protocol also differed—one expert had been involved in feasibility testing, whereas the other two were naïve to the protocol.

Raters received training materials, including video demonstrations and practice T2-weighted MRI scans, along with a virtual Zoom session led by an instructor. This session allowed them to review the materials, learn ITK-SNAP procedures, and ask questions. Raters then practiced segmenting 1-2 scans, which were reviewed in a follow-up meeting where the instructor provided feedback and approved their readiness. Reliability testing began only after instructor approval. Each training session lasted 1–2 hours. Inter-rater reliability was assessed among the 3 raters, in addition to intra-rater reliability following >2-week delay.

The reliability testing featured a collection of 24 MRI scans. Of these scans, 7 were neurodevelopmentally typical children (ages 4-15 years), 9 adults without dementia (ages 31-94 years), 5 older adults with mild cognitive impairment (ages 70-75 years), and 3 older adults with dementia (ages 76-79 years). These scans were sourced from either ADNI (clinical specimens as well as older, neurotypical adults, adni.loni.usc.edu) or HSG members. The ADNI was launched in 2003 as a public-private partnership, led by Principal Investigator Michael W. Weiner, MD. The primary goal of ADNI has been to test whether serial magnetic resonance imaging (MRI), positron emission tomography (PET), other biological markers, and clinical and neuropsychological assessment can be combined to measure the progression of mild cognitive impairment (MCI) and early Alzheimer’s disease (AD).

Due to the sample MRI scans coming from different research sites, either on Siemens or Philips MRI scanners depending on scan-site, the method used when obtaining the scans varied. The variability in specimen demographic details and acquisition does, however, boosts the generalizability of the results from the reliability testing as practices vary across research fields. Scan presentation order was randomized to reduce order effects, and raters were not given any information regarding the participants beyond the MRI scans themselves. All scans had 0.4x0.4x2 mm^3^ resolution obtained with 3T field strength. Scans were aligned perpendicular to the long axis of the hippocampus and in the coronal plane. Furthermore, all scans were selected through a quality control procedure based on the visibility of landmarks which were of key importance for the protocol, such as the SRLM (Canada et al., 2024). Imaging artifacts such as motion effects and reconstruction errors were included if mild or moderate, excluded if severe.

### 2.5. Statistical analyses

During the Delphi procedure, ratings were collected on 9-point Likert scales and summarized using descriptive statistics. Consensus was defined as a statistical majority, with responses recoded as clear (6–9) vs. unclear (1–5), and agreement (ratings 6–9) vs. non-agreement (1–5). Differences in frequency were evaluated using binomial tests (α = 0.05).

For the initial check for feasible reliability, inter-rater agreement on the ranging rule was evaluated using Cohen’s κ, while agreement was assessed through the calculation of Intraclass Correlation Coefficients (ICC(2,k); Shrout and Fleiss, 1979) and Dice Similarity Coefficient (DSC; Dice, 1945; Zou et al., 2004). For the formal reliability test, intra-rater reliability was assessed for absolute agreement (ICC(2,1)) and average DSC on a subset of 11 scans, with segmentations performed twice by the three raters, separated by at least two weeks. To assess the reliability of the harmonized protocol, inter-rater reliability was measured across 24 cases using intraclass correlation coefficients (ICC(2,k)) and average DSC across all rater pairs. Consistency of inter-rater DSC between child and adult age groups, and by cognitive diagnosis (from ADNI data), was tested using non-parametric Mann-Whitney U and Kruskal-Wallis tests, respectively.

## 3. Results

### 3.1. Feasible assessment of a reliability study of the proposed protocol

The initial feasibility assessment indicated that the protocol could be implemented reliably, as raters demonstrated substantial agreement on the ranging (Cohen’s κ = 0.71 ± 0.13) and excellent consistency (ICC(2) = 0.99) and spatial overlap (DSC = 0.91 ± 0.02) for HT segmentation.

When asked on a 9-point scale (0—Not at All, 8—Extremely Well), the raters indicated that they understood the protocol well (rater 1 = 7, rater 2 = 8) and that the rules were clearly defined in the training documents (rater 1 = 6, rater 2 = 8). When asked about the overall difficulty of performing the tracings on a 9-point scale (0—Very Easy, 8—Very Difficult), the raters indicated that the protocol was easy to apply (rater 1 = 2, rater 2 = 0). When asked more specifically about the different rules on a 7-point scale (0—Very Easy, 6—Very Difficult), the raters indicated that all the rules were easy to apply (Posterior: rater 1 = 1, rater 2 = 0; Inferior: rater 1 = 2, rater 2 = 0; Superior: rater 1 = 2, rater 2 = 0; Lateral: rater 1 = 2, rater 2 = 0), with the exception of the medial boundary which was considered “neutral - 3” by rater 1 (rater 2 = 0). The feasibility test results and qualitative reporting were provided as background information for the Delphi procedure panel.

### 3.2. Delphi procedure evaluating the boundaries

The Delphi procedure included responses from 21 participating laboratories: four with experience in segmenting individual subfields within the tail, and 17 that combined all subfields into a single ‘tail’ label. All laboratories had over five years of experience with hippocampal segmentation; 95% had expertise with hippocampal measures on 3T data, and 86% had experience with relevant T_2_-weighted images. One laboratory reported experience limited to histology. After a single iteration, all boundary definitions were deemed clear and reached consensus (binomial test p < 0.001 for all; Table 1). No revisions to the rule definitions were necessary to achieve consensus, although the working group incorporated feedback to refine wording and update sample images for clarity. Note that the online supplement reports all comments.

**Table 1:**
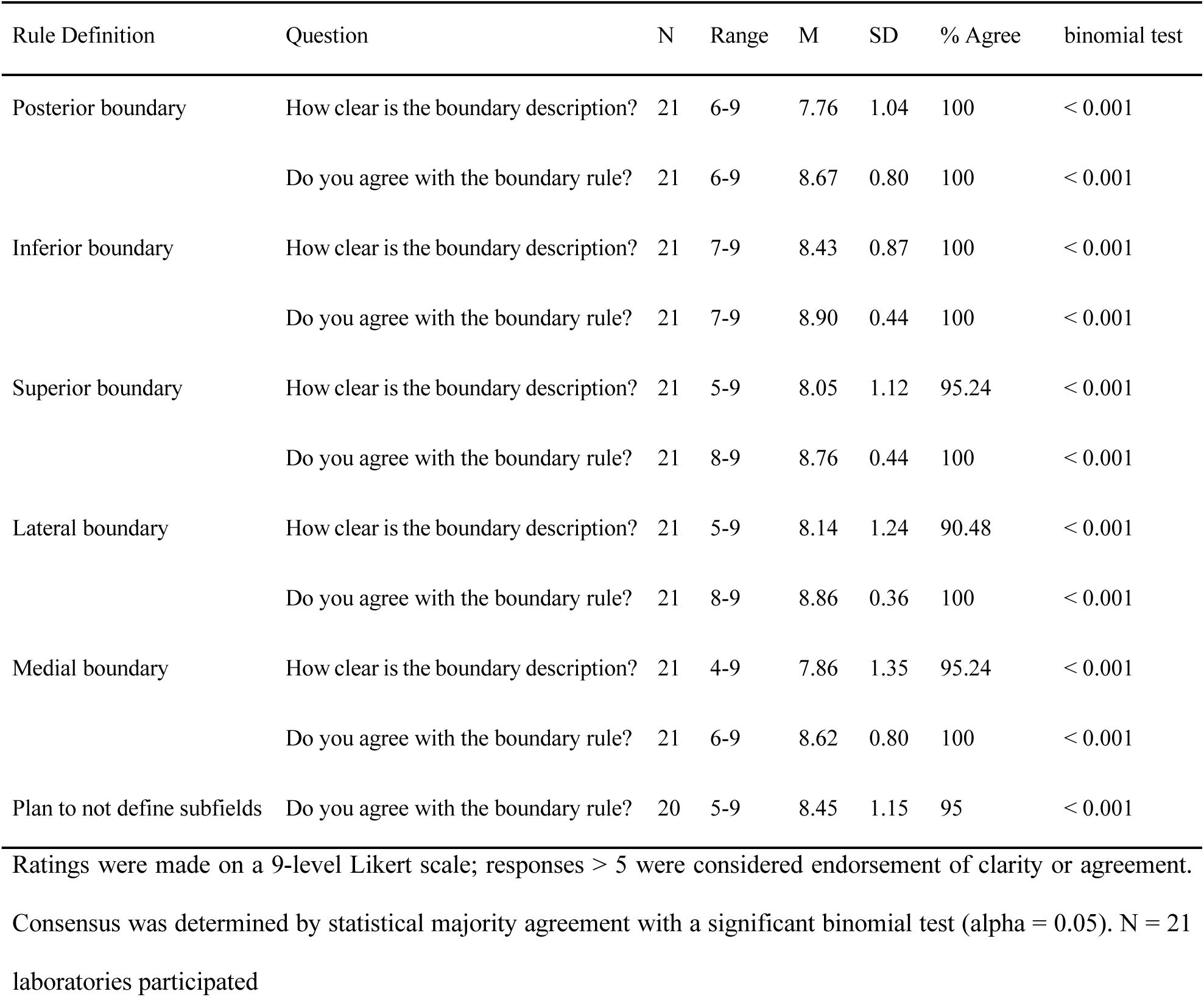
Summary of Delphi evaluation responses.

### 3.3. Formal reliability analysis of the consensus protocol

Intra-rater reliability, assessed by two experts and one novice after a delay of more than two weeks (n = 11), showed excellent agreement (Table 2). Consistency was observed among expert raters (ICC(2,1) = 0.85–1.00; DSC = 0.85–0.96) and the novice rater (ICC(2,1) = 0.92– 1.00; DSC = 0.81–0.96). Comparable reliability between experts and the novice indicates that the protocol can be applied consistently regardless of prior experience with manual segmentation or hippocampal anatomy. However, familiarity with the specific protocol may promote faster attainment of reliability. Inter-rater reliability was tested on the sample of N = 24 brains and found to be excellent for the left and right HT (ICC(2,k) = 0.95–0.99; DSC = 0.71–0.93, see Table 2).

**Table 2:** Summary intra- and inter-rater reliability of the harmonized protocol.

| Region | Intra-Rater |  | Inter-Rater |  |
| --- | --- | --- | --- | --- |
|  | (3 raters, n = 11) |  | (3 raters, n = 24) |  |
|  | Avg. *ICC*(2) | Avg. *DSC* | *ICC*(2,k) | Avg. *DSC* |
| HT-L | 0.99 | 0.92 ± 0.03 | 0.99 | 0.88 ± 0.04 |
| HT-R | 0.98 | 0.92 ± 0.03 | 0.98 | 0.86 ± 0.04 |

Reliability was tested on a blinded dataset by two expert raters and 1 novice rater (k = 3 raters), all naïve to the protocol. Average ICC, and average DSC (+ SD) are reported. ICC—intra-class correlation coefficient; DSC—Dice Similarity Coefficient; HT—Hippocampal Tail

Reliability was comparable across raters, independent of the demographic characteristics of the scanned participants. Within the limits of the current sample, these results offer preliminary evidence that reliability does not vary systematically by age group (children versus adults) or by cognitive status in older adults when applying ADNI procedures (Table 3).

**Table 3:** Summary of average inter-rater dice similarity coefficient by scan demographic.

| Region | Age-Group Comparisons |  |  | Comparisons by Cognitive Diagnosis |  |  |  |
| --- | --- | --- | --- | --- | --- | --- | --- |
|  | Children<br>(n=7) | Adults<br>(n=17) | p-value | Cognitive Typical<br>(n = 5) | MCI<br>(n = 5) | Dementia<br>(n=3) | p-value |
| HT-L | $0.89 \pm 0.02$ | $0.88 \pm 0.04$ | 0.58 | $0.88 \pm 0.04$ | $0.89 \pm 0.04$ | $0.87 \pm 0.03$ | 0.69 |
| HT-R | $0.88 \pm 0.03$ | $0.85 \pm 0.04$ | 0.13 | $0.83 \pm 0.03$ | $0.87 \pm 0.05$ | $0.86 \pm 0.03$ | 0.14 |

Average dice similarity coefficient (DSC) + SD is reported stratified by age group and cognitive diagnosis that was associated with the scan. Distributions were compared for statistically significant difference by Mann-Whitney U and Kruskal-Wallis tests ( $\alpha = 0.05$ ). Comparisons by cognitive diagnosis were made only among scans collected by ADNI and included standardized protocol for diagnosis. MCI—Mild cognitive impairment; HT—Hippocampal Tail.

### 3.4. HSG Harmonized Protocol for segmenting the hippocampal tail

Following community consensus and confirmation of reliability, we formalized the HSG Harmonized Protocol for segmentation of the HT. The HT is drawn as a single label, without distinguishing its subfields (Figure 3). The procedure begins by selecting the anterior-posterior range of the hippocampal tail. Note that the anterior HT slice refers to the slice immediately posterior to the final body slice, which is the most posterior slice on which the lamina quadrigemina (comprising superior and inferior colliculi) is clearly visible, as described in our previous publications (Olsen et al., 2019; Daugherty et al., 2025). Then, the HT is segmented by applying a set of four rules defining the outer boundaries of the structure. A summary of the complete, harmonized segmentation protocol for the HT is provided here; examples and training materials are available for download (https://hippocampalsubfields.com/harmonized-protocol/).

**Figure 3:**
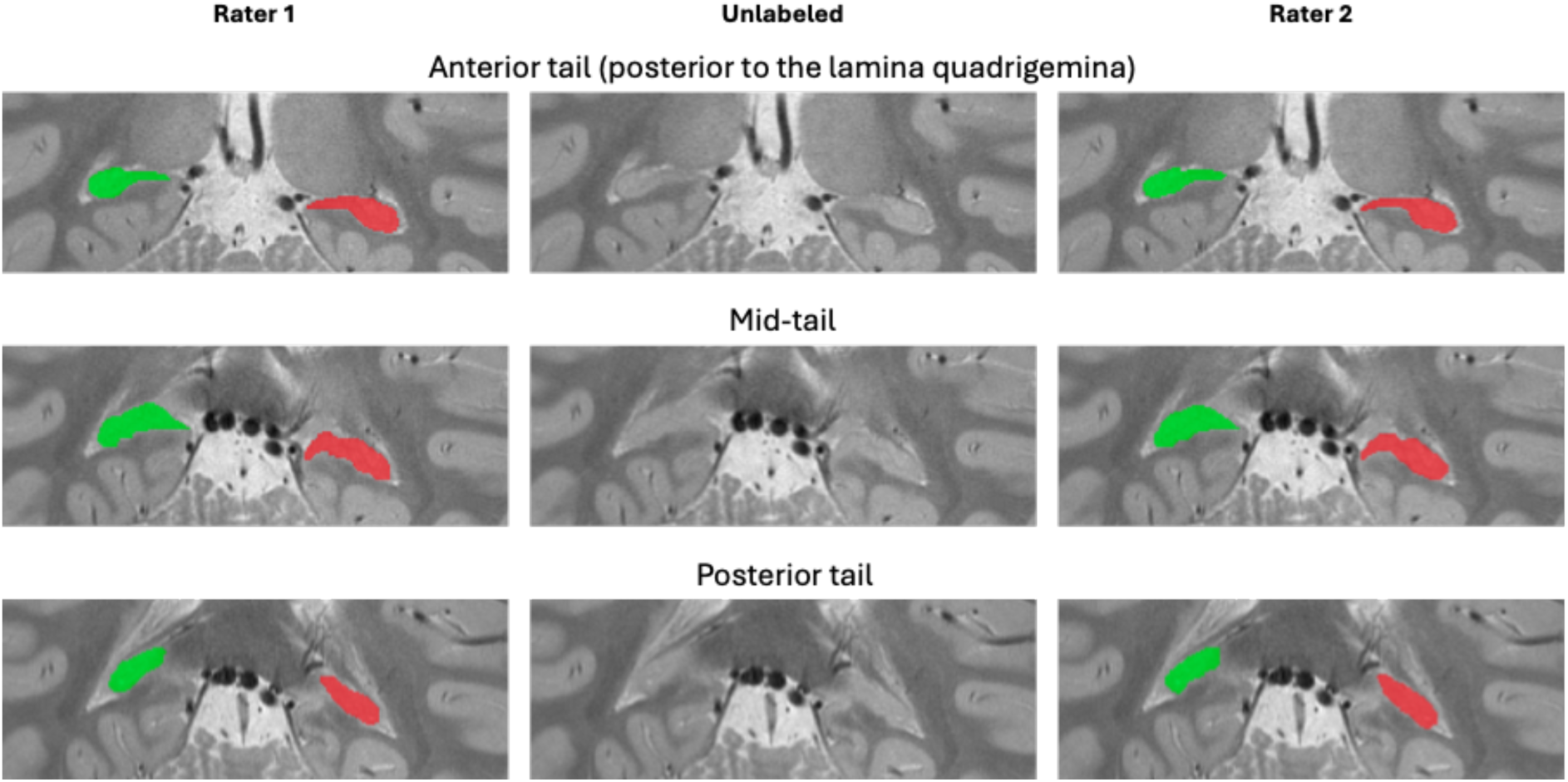
Segmentations on example MRI by two raters from the feasibility assessment. Green: right and red: left HT segmentation

#### 3.4.1. Posterior boundary

The HT is segmented only when it is clearly visible and distinguishable from partial volume voxels, which often result from bright CSF and are excluded from labeling. In some cases, sagittal slices should also be checked to identify the most posterior or lateral extent of the structure, particularly when partial volume effects occur (see Figure 4). The final HT slice may vary between hemispheres due to misaligned acquisitions or asymmetrical hippocampi. As outlined in the “medial border” section, the HT might detach from the subsplenial gyrus (Mai et al., 2008), and only the lateral (mostly oval shaped) part of the HT is segmented (see Figure 4 a-2).

**Figure 4:**
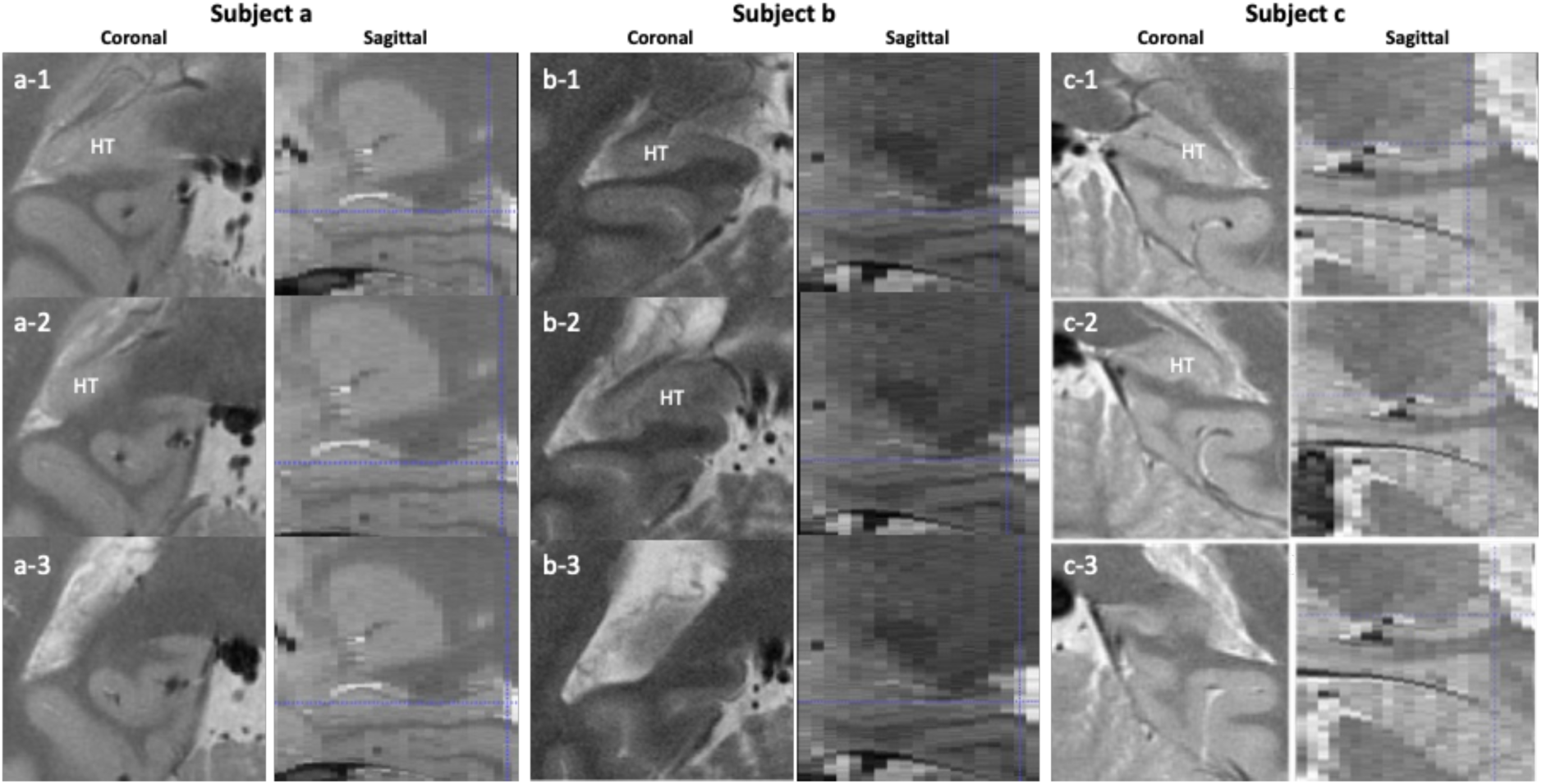
Posterior boundary. Three HT slices - anterior (1) to posterior (3) - shown for three different subjects (a, b and c) in both coronal and sagittal views. The last slices on which the HT should be segmented are slices a-2, b-2 and c-2. Partial voluming between HT (dark) and CSF (bright) can be seen in a-3 and b-3.

#### 3.4.2. Inferior boundary

The HT is bordered inferiorly by the white matter of the fusiform and lingual gyri as well as the cingulum bundle (Oishi et al., 2012). Consistent with the outer boundaries protocol of the hippocampal body (Daugherty et al., 2025), the inferior boundary of the HT is defined by the border between the gray matter of the hippocampus and the white matter located inferior to it, such that white matter is excluded from the definition of the HT (Figure 5).

**Figure 5:**
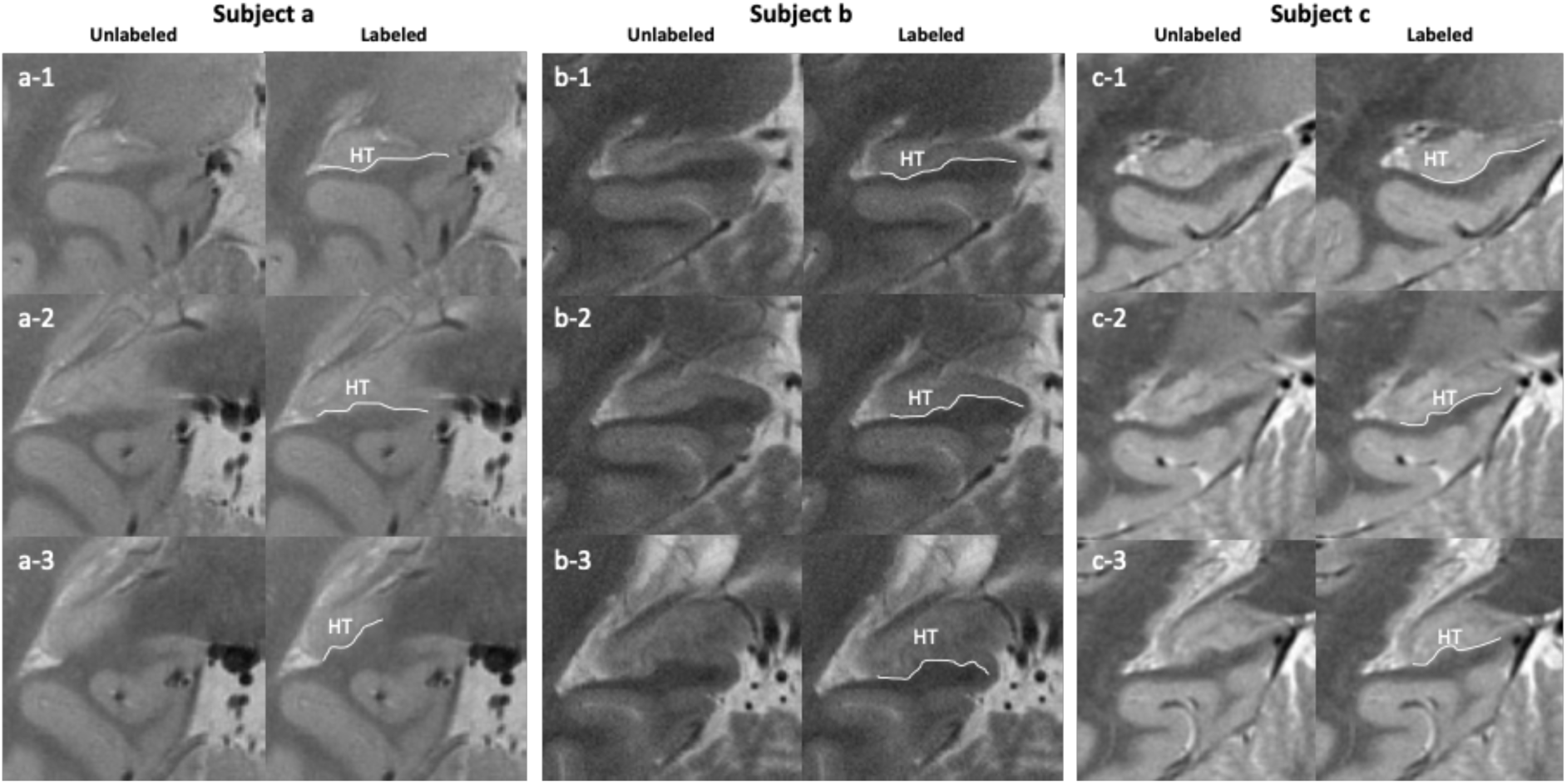
Inferior boundary. Three HT slices -anterior (1) to posterior (3) - shown for three different subjects (a, b and c).

#### 3.4.3. Superior boundary

The fimbria, the splenium of the corpus callosum, and/or the CSF constitute the superior border of HT (see Figure 6). For some cases, a small piece of gray matter appears to detach from the main portion of the HT (see the dotted line in Figure 6 b-2). This detached part should not be considered HT as it could belong to either hippocampal or thalamic tissue. If the gray matter is not detached, then the entire superior portion of the HT should be included. Note that the fimbria and the fornix should not be included in the segmentation.

**Figure 6:**
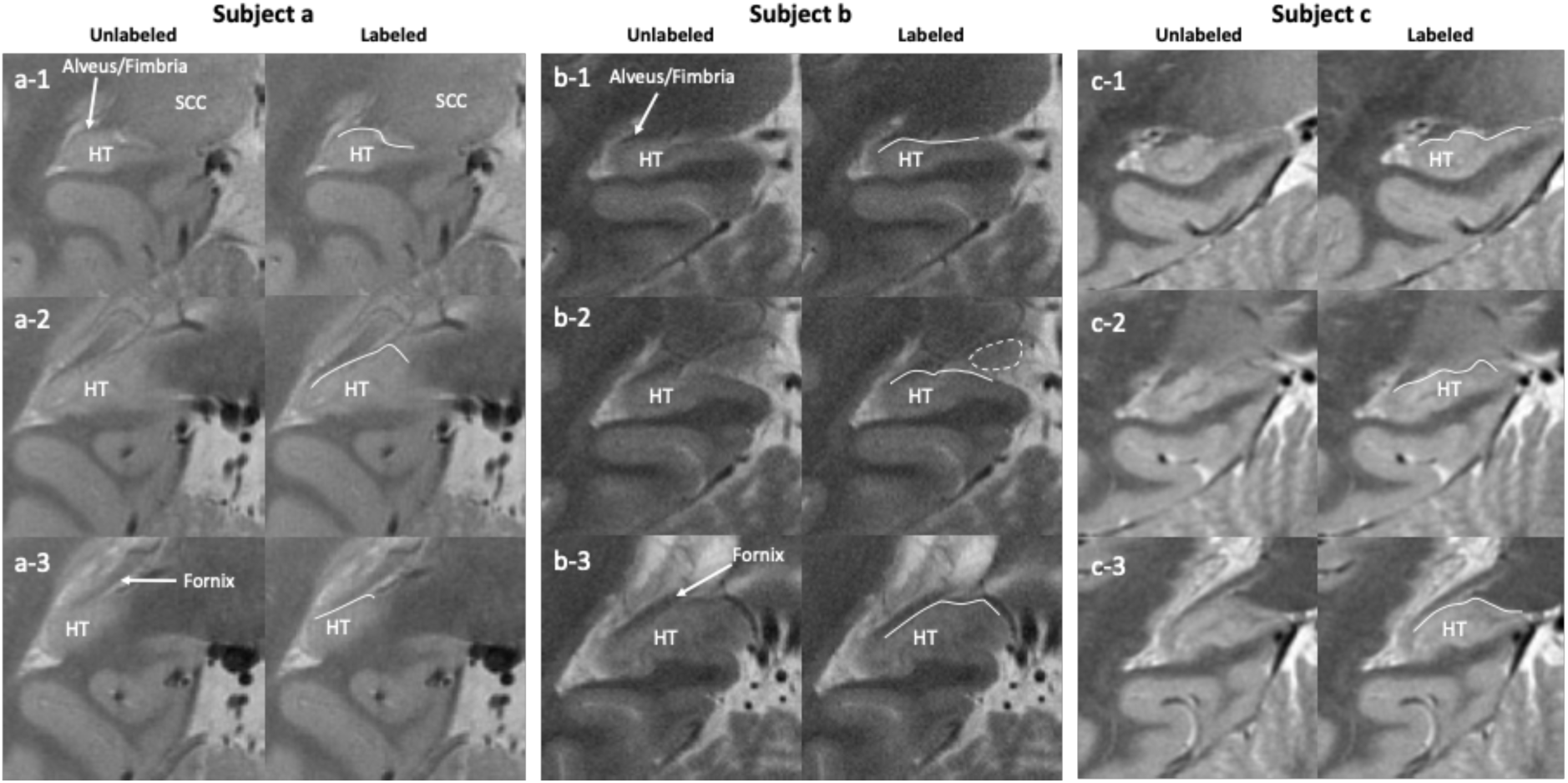
Superior boundary. Three HT slices - anterior (1) to posterior (3) - shown for three different subjects (a, b and c). The dotted line demarcates a detached part of gray matter that should not be considered HT (see text). SCC: splenium of the corpus callosum.

#### 3.4.5. Lateral boundary

The lateral boundary of the HT is defined by the alveus and/or CSF if the alveus is not visible, such that the alveus should not be included in the segmentation (Figure 7). Partial voluming may be observed in this region given the mixture of signals stemming from a variety of proximal tissues/structures (gray matter, white matter, CSF, and choroid plexus), and so should be considered as part of the HT. Consistent with the recommendations in the outer boundaries protocol for the hippocampal body (Olsen et al., 2019; Daugherty et al., 2025), contiguous slices should be consulted to evaluate whether the portion of tissue falls within the hippocampal region.

**Figure 7:**
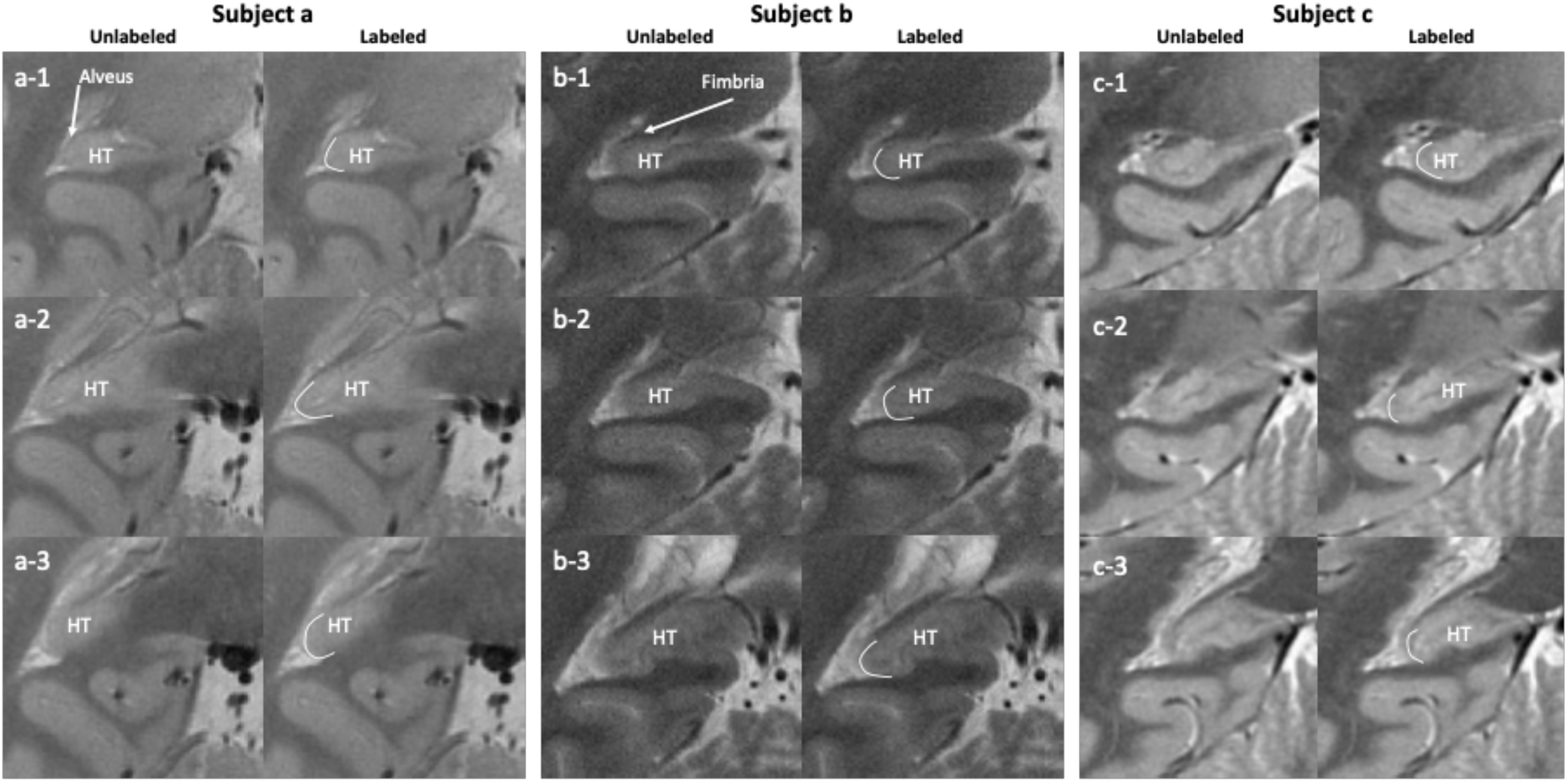
Lateral boundary. Three HT slices -anterior (1) to posterior (3) - shown for three different subjects (a, b and c).

#### 3.4.5. Medial boundary

Consistent with the outer boundaries protocol of the hippocampal body (Olsen et al., 2019; Daugherty et al., 2025), the medial boundary between the HT and parahippocampal cortex is defined as the most supero-medial corner of the parahippocampal gyrus (see the red dots in Figure 8 a-1, a-2, b-1, b-2, b-3, c-1, c-2, c-3). The boundary is placed at the level of maximum curvature in the cortical ribbon. The HT blends with a gyrus sometimes referred to as subsplenial gyrus (Duvernoy et al., 2013). Given that the subsplenial gyrus contains a mixture of CA1 and CA3, this gyrus is segmented together with the HT and considered the same structure, as long as these two regions are connected. When the HT detaches from the subsplenial gyrus (Mai et al., 2008), the medial portion is excluded and only the lateral (mostly oval shaped) part of the HT is segmented. More medially or when the HT is detached from the subsplenial gyrus (Figure 8 a-3), the boundary is defined by the border between the gray matter of the hippocampus and the white matter.

**Figure 8:**
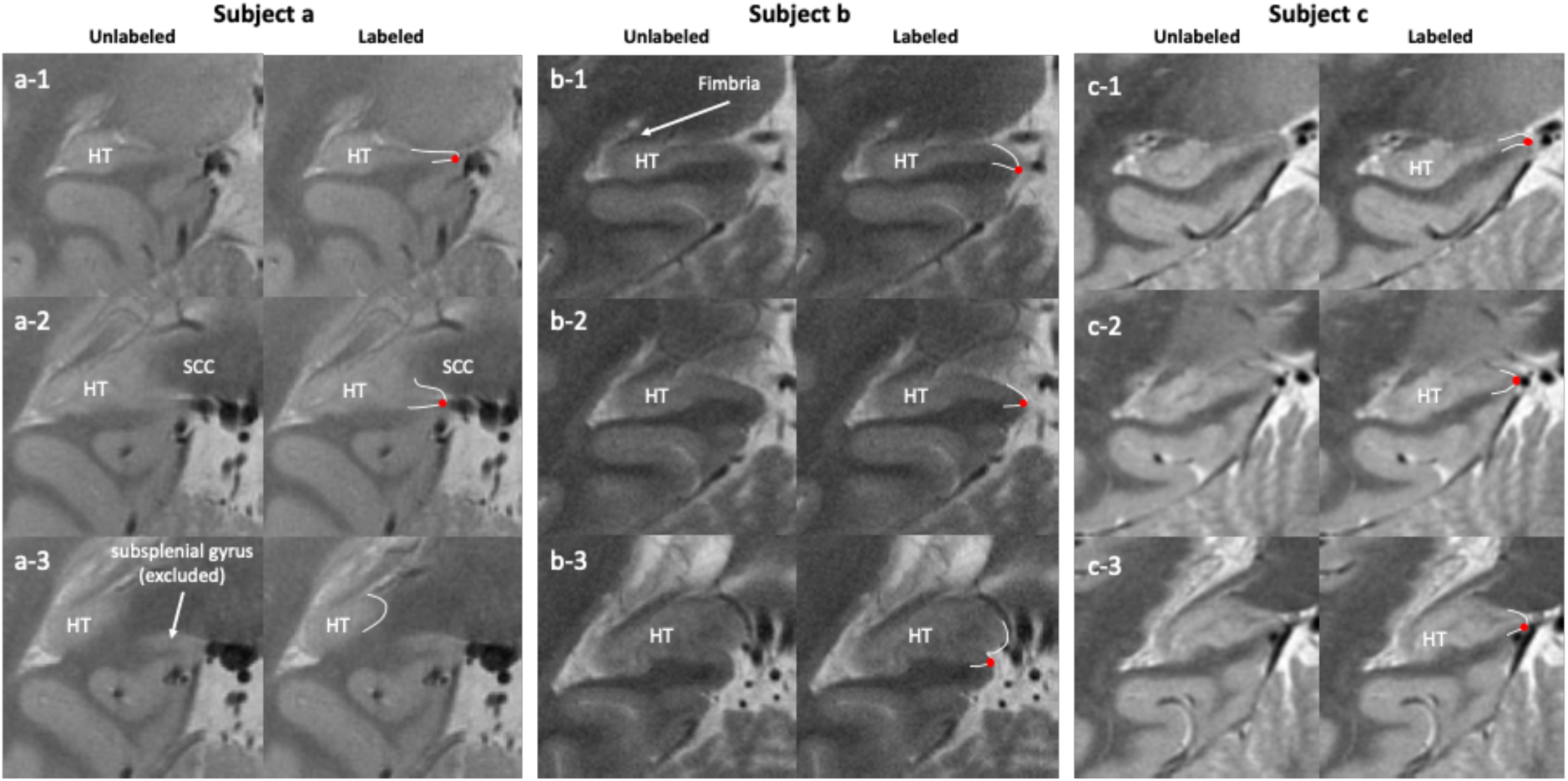
Medial boundary. Three HT slices - anterior (1) to posterior (3) - shown for three different subjects (a, b and c). For subject a, the HT detaches from the subsplenial gyrus (a-3). SCC: splenium of the corpus callosum. The red dots represent the most supero-medial corner of the parahippocampal gyrus.

## 4. Discussion

In the present manuscript, the HSG presents a harmonized protocol for segmentation of the HT on high-resolution *in vivo* MRI. This protocol is reliable and freely available on the HSG website (https://hippocampalsubfields.com/harmonized-protocol/). It complements the previously published harmonized protocol for subfield segmentation in the hippocampal body (Daugherty et al., 2025).

In the present protocol, we chose to consider the HT as a unitary region. This decision, supported by the Delphi procedure, was based on two key factors: first, subfield segmentation based on visible landmarks such as the SRLM is less feasible in the HT; second, and more importantly, the shape of the HT on coronal slices varies greatly depending on the orientation of the structure (Adler et al., 2018; de Flores et al., 2020). Using *ex vivo* data, de Flores et al. (2020) identified two shape patterns, with one resembling the hippocampal body. Notably, the non-body shape could appear similar to the body-like group when reoriented, as shown in Figure 2. Since the current HSG protocol is intended for use on commonly available anisotropic T_2_-weighted images (1–3 mm slice thickness), reorienting the scans without introducing considerable distortion is not feasible.

To encourage participation in the Delphi procedure, which was an essential part of harmonization, we issued open calls via listservs, our website, social media, and international conference presentations. We prioritized input from both experts in hippocampal subfield research and novices to improve accessibility. As in our prior work on HSG harmonized protocol boundary definitions (Olsen et al., 2019; Daugherty et al., 2025), the Delphi procedure achieved consensus in a single voting round. Qualitative feedback from feasibility raters and Delphi panelists informed iterative refinement of the protocol descriptions, supporting materials, and interpretation of the achieved consensus. All boundary definitions received high clarity and agreement, indicating strong community support. Notably, reliability metrics were comparable between experts and a novice rater, showing that prior segmentation experience is not required to apply the protocol. This aligns with our goal of broad usability. Additionally, hippocampal subfield DSC values were consistent across child and adult scans, and within ADNI data, regardless of cognitive status. However, while our analyses demonstrated reliability across demographic characteristics, the relatively small sample size highlights the need for validation in larger, more representative cohorts.

The protocol was designed for use with any modern software supporting manual segmentation; for accessibility, all example materials are provided in ITK-Snap (Yushkevich et al., 2006). There are no specific requirements for hardware or tracing environment—lab-specific variations are acceptable if reliability matches reported metrics. While standardizing equipment is impractical for harmonized measurements across multiple labs across the globe, equipment variations may introduce variability that remains unknown. We hope that broader adoption of this protocol will facilitate evaluation of software and hardware effects (e.g. tablet vs. mouse) in the future studies. We anticipate the HSG Harmonized Protocol for the HT will be applied to existing and new high-resolution T_2_-weighted datasets, serving as a reference to align published findings under a common nomenclature for improved comparability.

Several limitations of the protocol and continuing work should be noted. First, the HT protocol was developed in accord with the general imaging requirements established by the HSG, and applied to subfield segmentation of the head and body (Olsen et al., 2019; Daugherty et al., 2025). These requirements assume sub-millimeter in-plane coronal resolution with T_2_-weighting, as this level of detail is necessary for accurate identification of key anatomical landmarks in the head and body regions (Wisse et al., 2017). However, because the HT protocol does not require delineation of individual subfields, it is more flexible and can likely be applied to lower-resolution datasets, including 1 mm³ T1-weighted images. While such data do not provide sufficient detail for subfield-level segmentation in the head and body, the simpler anatomy of the tail makes its application feasible. Note that certain boundaries described in the present protocol, particularly the superior one, can be challenging to apply, and users should ensure tracer reliability in regions where gradients and partial voluming make segmentation more difficult. Looking forward, advances in MRI acquisition, particularly the growing availability of high-resolution isotropic scans, will allow image reslicing and greater anatomical precision, creating opportunities to incorporate subfield segmentation in the tail as well. These developments position the HSG to continue expanding and refining the protocol to keep pace with evolving imaging technologies.

Second, the outlined protocol applies specifically to the HT and is intended to complement the previously developed protocol for the hippocampal body (Olsen et al., 2019; Daugherty et al., 2025). Researchers can use both protocols together for more comprehensive segmentation across these regions while the HSG continues developing guidelines for the hippocampal head. Once the head protocol is finalized, all three components will provide a harmonized framework for consistent segmentation throughout the entire hippocampus. Until then, applying the tail protocol alongside the body protocol offers a practical interim solution for studies aiming to include these regions in quantitative analyses.

## 5. Conclusion

The HSG harmonized protocol for the hippocampal tail offers a reliable, accessible approach for consistent segmentation across labs and MRI scanners, and complements the existing HSG body protocol, with a head protocol forthcoming. By establishing standardized guidelines, the HSG aims to improve comparability of findings across multiple sites that are engaged in developmental, aging, and clinical research. While the current method is optimized for high-resolution T_2_-weighted images, its flexibility and compatibility with ongoing technological advances ensure its relevance and potential for future refinements.

## Supporting information

Supplement_Document_Tail

## Acknowledgements

In memory of Dr. Ricardo Insausti, a dear friend and colleague who shared his expertise and love of the hippocampus with this group.

This work was supported NIH/NIA R01-AG070592 (multi-site award; lead PI L. Wang); NIH/NIA P30-AG072931 (A.M. Daugherty); NIH/NIA R01-AG011230 (A.M. Daugherty and N. Raz); NIH/NICHD F32-HD108960 (K.L. Canada); NIH/NIA R01-AG073250 (T. Brown); NIH/NIMH R01-MH107512 (N. Ofen); NIH/NICHD R01-HD079518 (T. Riggins); NIH/NIA P30-AG066519 (C. Stark); NIH/NIA RF1-AG056014 and R01-AG069474 (P. Yushkevich); Fondation Alzheimer, Région Normandie, Fondation Recherche Alzheimer and Fondation Philippe Chatrier (all to R. de Flores); Canadian Institutes of Health Research (CIHR) PJT-162292 (R.K. Olsen); MultiPark -A Strategic Research Area at Lund University; Bente Rexed Gerstedt Foundation (both to L. Wisse).

Portions of the included MRI scans included in this project was funded by the Alzheimer’s Disease Neuroimaging Initiative (ADNI) (National Institutes of Health Grant U01 AG024904) and DOD ADNI (Department of Defense award number W81XWH-12-2-0012). ADNI is funded by the National Institute on Aging, the National Institute of Biomedical Imaging and Bioengineering, and through generous contributions from the following: AbbVie, Alzheimer’s Association; Alzheimer’s Drug Discovery Foundation; Araclon Biotech; BioClinica, Inc.; Biogen; Bristol-Myers Squibb Company; CereSpir, Inc.; Cogstate; Eisai Inc.; Elan Pharmaceuticals, Inc.; Eli Lilly and Company; EuroImmun; F. Hoffmann-La Roche Ltd and its affiliated company Genentech, Inc.; Fujirebio; GE Healthcare; IXICO Ltd.; Janssen Alzheimer Immunotherapy Research & Development, LLC.; Johnson & Johnson Pharmaceutical Research & Development LLC.; Lumosity; Lundbeck; Merck & Co., Inc.; Meso Scale Diagnostics, LLC.; NeuroRx Research; Neurotrack Technologies; Novartis Pharmaceuticals Corporation; Pfizer Inc.; Piramal Imaging; Servier; Takeda Pharmaceutical Company; and Transition Therapeutics. The Canadian Institutes of Health Research is providing funds to support ADNI clinical sites in Canada. Private sector contributions are facilitated by the Foundation for the National Institutes of Health (www.fnih.org). The grantee organization is the Northern California Institute for Research and Education, and the study is coordinated by the Alzheimer’s Therapeutic Research Institute at the University of Southern California. ADNI data are disseminated by the Laboratory for Neuro Imaging at the University of Southern California.

We would like to thank our former Advisory Council members Pr. Arne Ekstrom, Dr. Geoff Kerchner and Pr. Michael Yassa for their valuable contributions to the group’s work.

## Resource and Data Sharing Statement

All supporting data and resource materials are available for download from the Hippocampal Subfield Group (HSG) website: https://hippocampalsubfields.com/harmonized-protocol/.

## References

1. Adler DH, Wisse LEM, Ittyerah R, Pluta JB, Ding S-L, Xie L, Wang J, Kadivar S, Robinson JL, Schuck T, Trojanowski JQ, Grossman M, Detre JA, Elliott MA, Toledo JB, Liu W, Pickup S, Miller MI, Das SR, Wolk DA, Yushkevich PA. 2018. Characterizing the human hippocampus in aging and Alzheimer’s disease using a computational atlas derived from ex vivo MRI and histology. Proceedings of the National Academy of Sciences of the United States of America 115:4252–4257.

2. Aggleton JP. 2012. Multiple anatomical systems embedded within the primate medial temporal lobe: implications for hippocampal function. Neuroscience and biobehavioral reviews 36:1579–96.

3. Bartsch T, Wulff P. 2015. The hippocampus in aging and disease: From plasticity to vulnerability. Neuroscience 309:1–16.

4. Boccardi M, Bocchetta M, Apostolova LG, Barnes J, Bartzokis G, Corbetta G, DeCarli C, DeToledo-Morrell L, Firbank M, Ganzola R, Gerritsen L, Henneman W, Killiany RJ, Malykhin N, Pasqualetti P, Pruessner JC, Redolfi A, Robitaille N, Soininen H, Tolomeo D, Wang L, Watson C, Wolf H, Duvernoy H, Duchesne S, Jack CR, Frisoni GB, EADC-ADNI Working Group on the Harmonized Protocol for Manual Hippocampal Segmentation. 2015. Delphi definition of the EADC-ADNI Harmonized Protocol for hippocampal segmentation on magnetic resonance. Alzheimer’s & dementia : the journal of the Alzheimer’s Association 11:126–38.

5. Brunec IK, Robin J, Patai EZ, Ozubko JD, Javadi A-H, Barense MD, Spiers HJ, Moscovitch M. 2019. Cognitive mapping style relates to posterior-anterior hippocampal volume ratio. Hippocampus 29:748–754.

6. Buckner RL. 2010. The Role of the Hippocampus in Prediction and Imagination. Annual Review of Psychology 61:27–48.

7. Canada KL, Mazloum-Farzaghi N, Rådman G, Adams JN, Bakker A, Baumeister H, Berron D, Bocchetta M, Carr VA, Dalton MA, de Flores R, Keresztes A, La Joie R, Mueller SG, Raz N, Santini T, Shaw T, Stark CEL, Tran TT, Wang L, Wisse LEM, Wuestefeld A, Yushkevich PA, Olsen RK, Daugherty AM, Group the HS. 2024. A (sub)field guide to quality control in hippocampal subfield segmentation on high-resolution T2-weighted MRI. Human Brain Mapping 45:e70004.

8. Carr V a, Rissman J, Wagner AD. 2010. Imaging the human medial temporal lobe with high-resolution fMRI. Neuron 65:298–308.

9. Chauveau L, Kuhn E, Palix C, Felisatti F, Ourry V, de La Sayette V, Chételat G, de Flores R. 2021. Medial Temporal Lobe Subregional Atrophy in Aging and Alzheimer’s Disease: A Longitudinal Study. Front Aging Neurosci 13:750154.

10. Chen KHM, Chuah LYM, Sim SKY, Chee MWL. 2010. Hippocampal region-specific contributions to memory performance in normal elderly. Brain and Cognition 72:400–407.

11. Daugherty AM, Carr V, Canada KL, Rådman G, Brown T, Augustinack J, Amunts K, Bakker A, Berron D, Burggren A, Chetelat G, Flores R de, Ding S-L, Huang Y, Insausti R, Johnson E, Kanel P, Keresztes A, Kedo O, Kennedy KM, Lee J, Malykhin N, Martinez A, Mueller S, Mulligan E, Ofen N, Palombo D, Pasquini L, Pluta J, Raz N, Riggins T, Rodrigue KM, Saifullah S, Schlichting ML, Stark C, Wang L, Yushkevich P, Joie RL, Wisse L, Olsen R, Initiative the ADN. 2025. Harmonized Protocol for Subfield Segmentation in the Hippocampal Body on High-Resolution in vivo MRI from the Hippocampal Subfields Group (HSG). :2025.04.29.651039. Available from: https://www.biorxiv.org/content/10.1101/2025.04.29.651039v1

12. Ding S-L. 2013. Comparative anatomy of the prosubiculum, subiculum, presubiculum, postsubiculum, and parasubiculum in human, monkey, and rodent. The Journal of comparative neurology 521:4145–62.

13. Duvernoy HM, Cattin F, Risold P-Y. 2013. The Human Hippocampus: Functional Anatomy, Vascularization and Serial Sections with MRI. Berlin, Heidelberg: Springer. Available from: https://link.springer.com/10.1007/978-3-642-33603-4

14. Fenton AA. 2024. Remapping revisited: how the hippocampus represents different spaces. Nat Rev Neurosci 25:428–448.

15. de Flores R, Berron D, Ding S-L, Ittyerah R, Pluta JB, Xie L, Adler DH, Robinson JL, Schuck T, Trojanowski JQ, Grossman M, Liu W, Pickup S, Das SR, Wolk DA, Yushkevich PA, Wisse LEM. 2020. Characterization of hippocampal subfields using ex vivo MRI and histology data: Lessons for in vivo segmentation. Hippocampus 30:545–564.

16. de Flores R, La Joie R, Chételat G. 2015. Structural imaging of hippocampal subfields in healthy aging and Alzheimer’s disease. Neuroscience 309:29–50.

17. Genon S, Bernhardt BC, La Joie R, Amunts K, Eickhoff SB. 2021. The many dimensions of human hippocampal organization and (dys)function. Trends Neurosci 44:977–989.

18. Geuze E, Vermetten E, Bremner JD. 2005. MR-based in vivo hippocampal volumetrics: 2. Findings in neuropsychiatric disorders. Molecular psychiatry 10:160–84.

19. Grande X, Wisse L, Berron D. 2023. Chapter 16 -Ultra-high field imaging of the human medial temporal lobe. In: Markenroth Bloch K, Guye M, Poser BA, editors. Advances in Magnetic Resonance Technology and Applications. Vol. 10. Ultra-High Field Neuro MRI. Academic Press. p 259–272. Available from: https://www.sciencedirect.com/science/article/pii/B9780323998987000316

20. Insausti R, Amaral DG. 2012. Hippocampal Formation. In: Mai J, Paxinos G, editors. The Human Nervous System. Third Edit. San Diego: Elsevier Academic Press. p 896–942.

21. Kalpouzos G, Chételat G, Baron J-C, Landeau B, Mevel K, Godeau C, Barré L, Constans J-M, Viader F, Eustache F, Desgranges B. 2009. Voxel-based mapping of brain gray matter volume and glucose metabolism profiles in normal aging. Neurobiology of aging 30:112–24.

22. Kim H. 2015. Encoding and retrieval along the long axis of the hippocampus and their relationships with dorsal attention and default mode networks: The HERNET model. Hippocampus 25:500–510.

23. Kolibius LD, Josselyn SA, Hanslmayr S. 2025. On the origin of memory neurons in the human hippocampus. Trends Cogn Sci 29:421–433.

24. Lepage M, Habib R, Tulving E. 1998. Hippocampal PET activations of memory encoding and retrieval: the HIPER model. Hippocampus 8:313–22.

25. Mai JK, Majtanik M, Paxinos G. 2008. Atlas of the human brain. New York, USA: Academic Press, Elsevier, New York, USA.

26. Maruszak A, Thuret S. 2014. Why looking at the whole hippocampus is not enough-a critical role for anteroposterior axis, subfield and activation analyses to enhance predictive value of hippocampal changes for Alzheimer’s disease diagnosis. Frontiers in cellular neuroscience 8:95.

27. Moscovitch M, Cabeza R, Winocur G, Nadel L. 2016. Episodic Memory and Beyond: The Hippocampus and Neocortex in Transformation. Annu Rev Psychol 67:105–134.

28. Moser EI, Kropff E, Moser M-B. 2008. Place Cells, Grid Cells, and the Brain’s Spatial Representation System. Annual Review of Neuroscience 31:69–89.

29. Mueller SG, Stables L, Du AT, Schuff N, Truran D, Cashdollar N, Weiner MW. 2007. Measurement of hippocampal subfields and age-related changes with high resolution MRI at 4T. Neurobiology of aging 28:719–26.

30. Oishi K, Faria AV, van Zijl PCM, Susumu Mori. 2012. MRI Atlas of Human White Matter. 2nd ed. Academic Press.

31. Olsen RK, Carr VA, Daugherty AM, La Joie R, Amaral RSC, Amunts K, Augustinack JC, Bakker A, Bender AR, Berron D, Boccardi M, Bocchetta M, Burggren AC, Chakravarty MM, Chételat G, de Flores R, DeKraker J, Ding S-L, Geerlings MI, Huang Y, Insausti R, Johnson EG, Kanel P, Kedo O, Kennedy KM, Keresztes A, Lee JK, Lindenberger U, Mueller SG, Mulligan EM, Ofen N, Palombo DJ, Pasquini L, Pluta J, Raz N, Rodrigue KM, Schlichting ML, Lee Shing Y, Stark CEL, Steve TA, Suthana NA, Wang L, Werkle-Bergner M, Yushkevich PA, Yu Q, Wisse LEM. 2019. Progress update from the hippocampal subfields group. Alzheimer’s & Dementia: Diagnosis, Assessment & Disease Monitoring 11:439–449.

32. Palomero-Gallagher N, Kedo O, Mohlberg H, Zilles K, Amunts K. 2020. Multimodal mapping and analysis of the cyto- and receptorarchitecture of the human hippocampus. Brain Struct Funct 225:881–907.

33. Plachti A, Eickhoff SB, Hoffstaedter F, Patil KR, Laird AR, Fox PT, Amunts K, Genon S. 2019. Multimodal Parcellations and Extensive Behavioral Profiling Tackling the Hippocampus Gradient. Cerebral Cortex 29:4595–4612.

34. Poppenk J, Evensmoen HR, Moscovitch M, Nadel L. 2013. Long-axis specialization of the human hippocampus. Trends in cognitive sciences 17:230–40.

35. Shrout PE, Fleiss JL. 1979. Intraclass correlations: uses in assessing rater reliability. Psychol Bull 86:420–428.

36. Small S a, Schobel S a, Buxton RB, Witter MP, Barnes C a. 2011. A pathophysiological framework of hippocampal dysfunction in ageing and disease. Nature reviews Neuroscience 12:585–601.

37. Strange BA, Witter MP, Lein ES, Moser EI. 2014. Functional organization of the hippocampal longitudinal axis. Nat Rev Neurosci 15:655–669.

38. Sun Y, Hu N, Wang M, Lu L, Luo C, Tang B, Yao C, Sweeney JA, Gong Q, Qiu C, Lui S. 2023. Hippocampal subfield alterations in schizophrenia and major depressive disorder: a systematic review and network meta-analysis of anatomic MRI studies. J Psychiatry Neurosci 48:E34–E49.

39. Ta AT, Huang SE, Chiu MJ, Hua MS, Tseng WYI, Chen SHA, Qiu A. 2012. Age-related vulnerabilities along the hippocampal longitudinal axis. Human Brain Mapping 33:2415–2427.

40. West MJ, Gundersen HJ. 1990. Unbiased stereological estimation of the number of neurons in the human hippocampus. The Journal of comparative neurology 296:1–22.

41. Wisse LEM, Daugherty AM, Olsen RK, Berron D, Carr VA, Stark CEL, Amaral RSC, Amunts K, Augustinack JC, Bender AR, Bernstein JD, Boccardi M, Bocchetta M, Burggren A, Chakravarty MM, Chupin M, Ekstrom A, de Flores R, Insausti R, Kanel P, Kedo O, Kennedy KM, Kerchner GA, LaRocque KF, Liu X, Maass A, Malykhin N, Mueller SG, Ofen N, Palombo DJ, Parekh MB, Pluta JB, Pruessner JC, Raz N, Rodrigue KM, Schoemaker D, Shafer AT, Steve TA, Suthana N, Wang L, Winterburn JL, Yassa MA, Yushkevich PA, la Joie R. 2017. A harmonized segmentation protocol for hippocampal and parahippocampal subregions: Why do we need one and what are the key goals? Hippocampus 27:3–11.

42. Wisse LEM, Wuestefeld A, Murray ME, Jagust W, La Joie R. 2025. Role of tau versus TDP-43 pathology on medial temporal lobe atrophy in aging and Alzheimer’s disease. Alzheimers Dement 21:e14582.

43. Wuestefeld A, Baumeister H, Adams JN, de Flores R, Hodgetts CJ, Mazloum-Farzaghi N, Olsen RK, Puliyadi V, Tran TT, Bakker A, Canada KL, Dalton MA, Daugherty AM, La Joie R, Wang L, Bedard ML, Buendia E, Chung E, Denning A, del Mar Arroyo-Jiménez M, Artacho-Pérula E, Irwin DJ, Ittyerah R, Lee EB, Lim S, del Pilar Marcos-Rabal M, Iñiguez de Onzoño Martin MM, Lopez MM, de la Rosa Prieto C, Schuck T, Trotman W, Vela A, Yushkevich P, Amunts K, Augustinack JC, Ding S-L, Insausti R, Kedo O, Berron D, Wisse LEM. 2024. Comparison of histological delineations of medial temporal lobe cortices by four independent neuroanatomy laboratories. Hippocampus 34:241–260.

44. Yushkevich PA, Amaral RSC, Augustinack JC, Bender AR, Bernstein JD, Boccardi M, Bocchetta M, Burggren AC, Carr VA, Chakravarty MM, Chételat G, Daugherty AM, Davachi L, Ding S-L, Ekstrom A, Geerlings MI, Hassan A, Huang Y, Iglesias JE, La Joie R, Kerchner GA, LaRocque KF, Libby LA, Malykhin N, Mueller SG, Olsen RK, Palombo DJ, Parekh MB, Pluta JB, Preston AR, Pruessner JC, Ranganath C, Raz N, Schlichting ML, Schoemaker D, Singh S, Stark CEL, Suthana N, Tompary A, Turowski MM, Van Leemput K, Wagner AD, Wang L, Winterburn JL, Wisse LEM, Yassa MA, Zeineh MM. 2015. Quantitative comparison of 21 protocols for labeling hippocampal subfields and parahippocampal subregions in in vivo MRI: Towards a harmonized segmentation protocol. NeuroImage [Internet]. Available from: http://linkinghub.elsevier.com/retrieve/pii/S1053811915000075

45. Yushkevich PA, Ittyerah R, Li Y, Denning AE, Sadeghpour N, Lim S, McGrew E, Xie L, DeFlores R, Brown CA, Wisse LEM, Wolk DA, Das SR, Alzheimer’s Disease Neuroimaging Initiative. 2024. Morphometry of medial temporal lobe subregions using high-resolution T2-weighted MRI in ADNI3: Why, how, and what’s next? Alzheimers Dement 20:8113–8128.

46. Yushkevich PA, Piven J, Hazlett HC, Smith RG, Ho S, Gee JC, Gerig G. 2006. User-guided 3D active contour segmentation of anatomical structures: Significantly improved efficiency and reliability. NeuroImage 31:1116–1128.

47. Zilioli A, Pancaldi B, Baumeister H, Busi G, Misirocchi F, Mutti C, Florindo I, Morelli N, Mohanty R, Berron D, Westman E, Spallazzi M. 2025. Unveiling the hippocampal subfield changes across the Alzheimer’s disease continuum: a systematic review of neuroimaging studies. Brain Imaging Behav 19:253–267.

48. Zou KH, Warfield SK, Bharatha A, Tempany CMC, Kaus MR, Haker SJ, Wells WM, Jolesz FA, Kikinis R. 2004. Statistical validation of image segmentation quality based on a spatial overlap index. Acad Radiol 11:178–189.

