## Supplement_Document_Tail for "Harmonized Protocol for Segmentation of the Hippocampal Tail on High-Resolution *in vivo* MRI from the Hippocampal Subfields Group (HSG)"

### Proposed Protocol for the Tail

#### On High-Resolution T2-weighted Images Collected at 3 Tesla

On behalf of the Tail Working Group

hippocampalsubfields.com

Acknowledgments and working group members: Robin de Flores, Kelsey Canada, Nicole Gervais, Anne Maass, Eóin Molloy, Gustaf Rådman and Jonathan Shine.

This document provides supplemental information for the evaluation of the proposed draft protocol to segment the hippocampal tail. Please do not implement this information in your own research or disseminate outside of the HSG until the protocol is evaluated and published.

#### Preface

This document describes the procedure and results of the feasibility assessment that was performed on a set of potential "rules" for ranging and outer boundary definitions to deliniate the hippocampal tail on high-resolution MRI.

Please review the supporting documentation included in this supplement alongside the online questionnaire to provide feedback on the procedures.

***Feasibility Assessment.*** The protocol was developed with the intent that it could be applied to brains imaged from different populations (e.g., children, healthy aging, Alzheimer's disease, epilepsy). A consideration was also made for typical imaging artifacts (e.g., motion, poor gray-white matter contrast) and variability in morphometry (e.g., round hippocampus shape vs. canonical shape) that occurs between persons, possibly in correlation with development or disease progression, and along the anterior-posterior axis. All of these sources of variability were represented in the feasibility data set.

Two expert raters participated in the feasibility assessment. One was part of the tail working group and the other was completely naive to the protocol prior to training. Training included detailed documentation with example image tracings, a 1-hour introductory training session (via Zoom), followed by prescribed practice and then an additional 1-hour of individualized feedback (via Zoom).

Raters showed substantial agreement on the ranging with Cohen's kappa = 0.71 ± 0.13 and had excellent agreement and overlap for segmenting the tail (N = 4 scans).

| **Rater 1, Average Volume Across Cases ($mm^3$; M $\pm$ SD)** | **Rater 2, Average Volume Across Cases ($mm^3$; M $\pm$ SD)** | **ICC(2)** | **Average Dice (M $\pm$ SD)** |
|---|---|---|---|
| 470.74 $\pm$ 155.41 | 473.86 $\pm$ 144.73 | 0.99 | 0.91 $\pm$ 0.02 |

All segmentations were completed throughout the length of the hippocampal tail, Example segmentations on three slices of the hippocampal tail on one MRI scan are shown. Please download the example segmentation files to review in ITK Snap for further detail.

Fig 1. Segmentations on example MRI by two raters from the feasibility assessment.

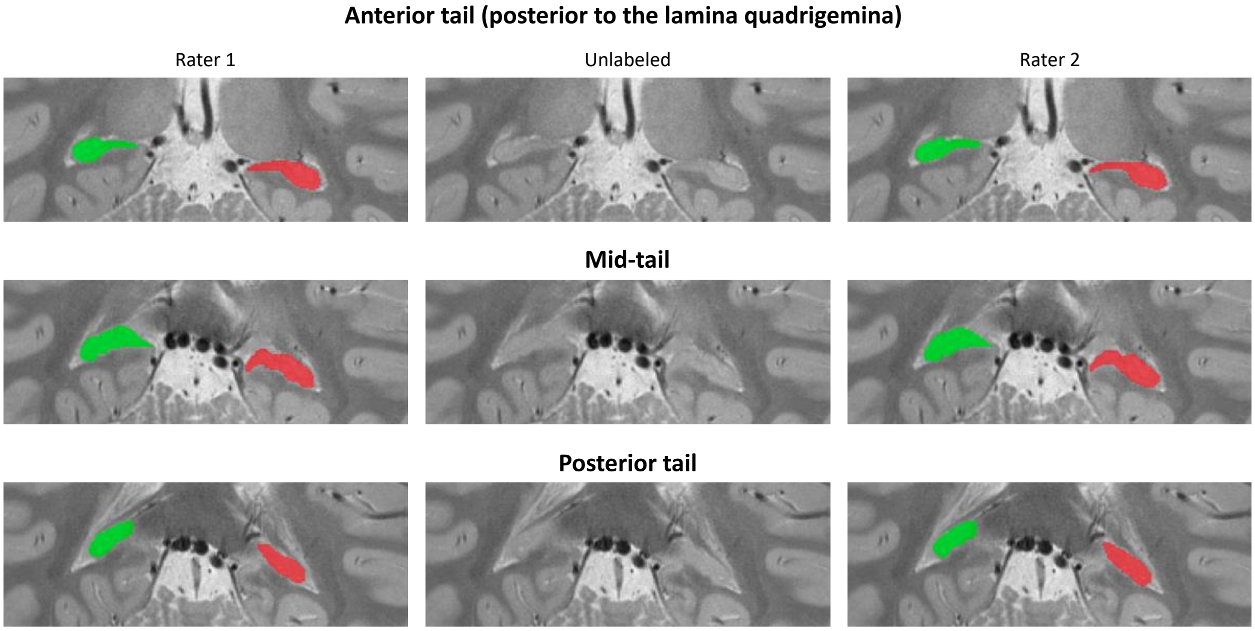


***Rater Experience.*** When asked on a 9-point scale (0—Not at All, 8—Extremely Well), the raters indicated that they understood the protocol well (rater 1 = 7, rater 2 = 8) and that the rules were clearly defined in the training documents (rater 1 = 6, rater 2 = 8). When asked about the overall difficulty of performing the tracings on a 9-point scale (0—Very Easy, 8—Very Difficult), the raters indicated that the protocol was easy to apply (rater 1 = 2, rater 2 = 0). When asked more specifically about the different rules on a 7-point scale (0—Very Easy, 6—Very Difficult), the raters indicated that all the rules were easy to apply (Anterior: rater 1 = 0, rater 2 = 0; Posterior: rater 1 = 1, rater 2 = 0; Ventral: rater 1 = 2, rater 2 = 0; Dorsal: rater 1 = 2, rater 2 = 0; Lateral: rater 1 = 2, rater 2 = 0), with the exception of the medial boundary which was considered "neutral - 3" by rater 1 (rater 2 = 0).

### Appendix A: Protocol Training Documentation



**Hippocampal Body Subfield Inner Boundary Protocol—Reliability Training Documentation**

The protocol was designed to be implemented in any software package that supports manual segmentation. For the purpose of the training and initial reliability assessment, we will be using ITKSnap.

To download ITKSnap: http://www.itksnap.org/pmwiki/pmwiki.php (last accessed 10/09/18)

**ITKSnap: Review of Relevant Tools**

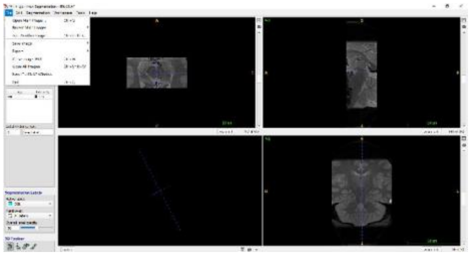

1. To load an image set: go to "Open New Image" under the **File** drop down menu
    a. Load either the DICOM series or NIFTI
    b. Click Finish
2. Create Segmentation Labels by clicking the pallet icon
    a. Load the Hippocampal Subfield Label.txt template by choosing "**Import Label Description**" under **Actions**

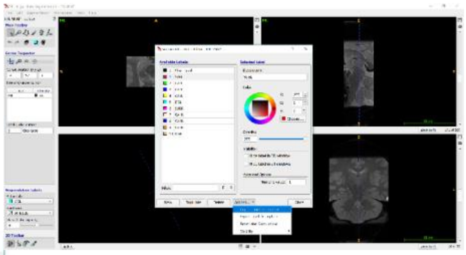

3. To Save Segmentation and Export Statistics go to **Segmentation** drop down menu
    a. Label the segmentation file as ID_your initial_date
    b. Export the statistics labeled as ID_your initial_stats

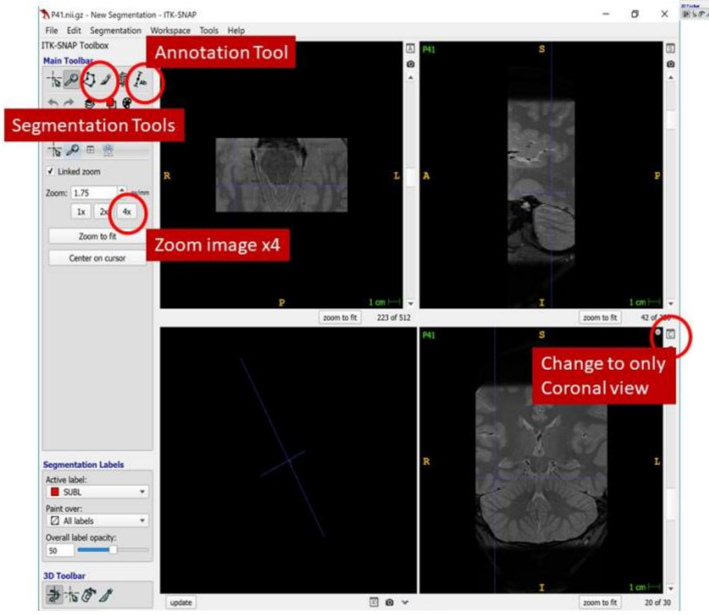


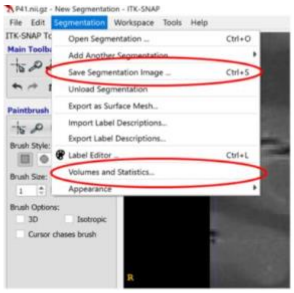

### Tail protocol

Robin de Flores, Nicole Gervais, Anne Maass and Jonathan Shine
On behalf of the Boundary Working Group

Last updated: November 2023

---

**Table of contents**

I. Anterior
II. Posterior
III. Ventral
IV. Dorsal
V. Lateral
VI. Medial

---

### 1. Anterior

Segmentation of the hippocampal tail (HT) begins on the slice where the colliculi (also called the lamina quadrigemina) are no longer visible (see Figure 1 a-2 & 1 b-2). Slices where the colliculi are partially visible are considered hippocampal body (HB) (Figure 1 b-1). The anterior boundary should be determined per-hemisphere. Specifically, due to the participant's position in the scanner, the hemispheres may appear misaligned in the resulting anatomical images. Figure 1b shows an example of slightly misaligned hemispheres. In case of severe misalignment (i.e. the colliculus is visible for one hemisphere but completely absent for the other), the anterior-most slice of the HT may differ between the left and right sides (please note that this is not the case for Figure 1b). In other words, only one colliculus (lower or upper) in a hemisphere needs to be visible for the slice to be considered HB in that hemisphere.

**Figure 1: Anterior boundary**
Anterior tail boundary for two example subjects, a) with aligned colliculi (left Column), b) with slightly misaligned colliculi (right column). Top row: Final posterior slice of the hippocampal body displaying the colliculi. Bottom row: Colliculi are no longer visible, and therefore this image is considered the first slice of the hippocampal tail.

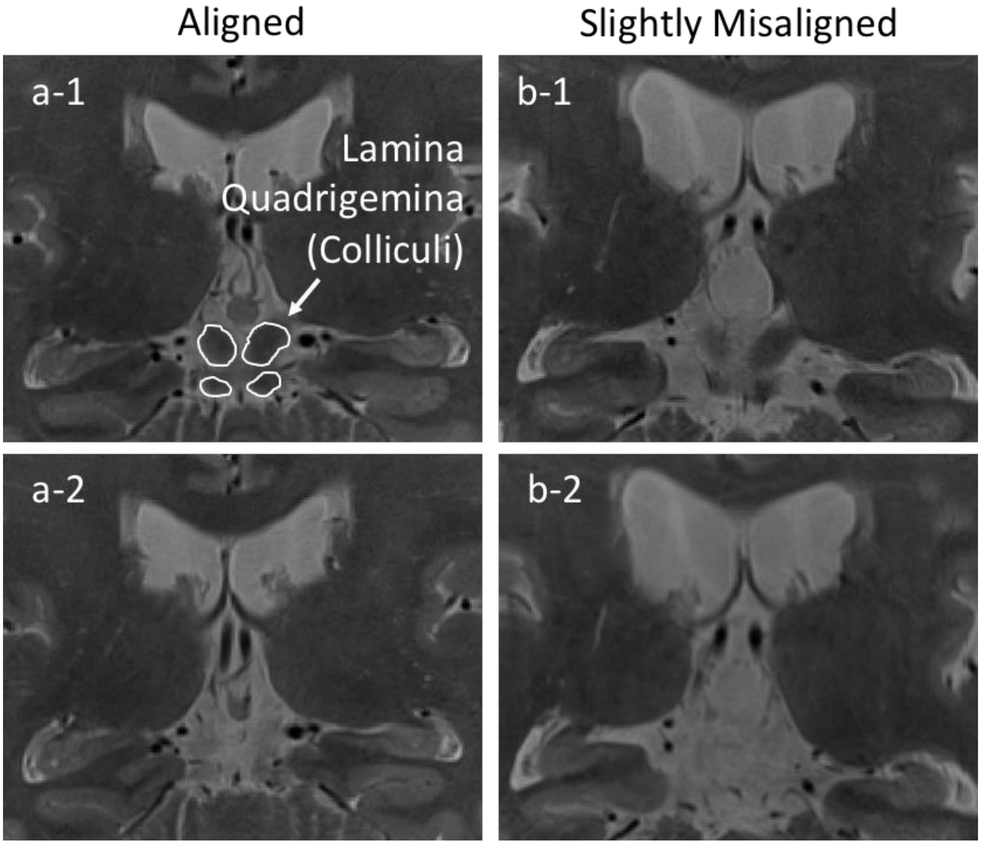

#### 2. Posterior

The HT is segmented as long as it is clearly visible (Figure 2; a1c, a2c and b1c, b2c). Partial volume voxels, often resulting from cerebrospinal fluid (CSF) which appears bright, are excluded from the label. It may be necessary to check for the HT also on sagittal slices to determine the most posterior/lateral extent of the structure, for instance if partial voluming occurs (Figure 2 a-3s and b-3s). The last HT slice may differ between hemispheres in case of misaligned acquisition or asymmetrical hippocampi. As explained in the "medial border" section, the HT might detach from the subsplenial gyrus (Mai et al., 2008) (Figure 2 a-2, a-3; Figure 6 a-3), and only the lateral (mostly oval shaped) part of the HT is segmented.

**Figure 2: Posterior boundary**
Three HT slices - anterior (1) to posterior (3) - shown for two different subjects (a and b) in both coronal "c" and sagittal "s" views. The last slices on which the HT should be segmented are slices a-2 and b-2. Partial voluming between HT (dark) and CSF (bright) can be seen in a-3c and b-3c.

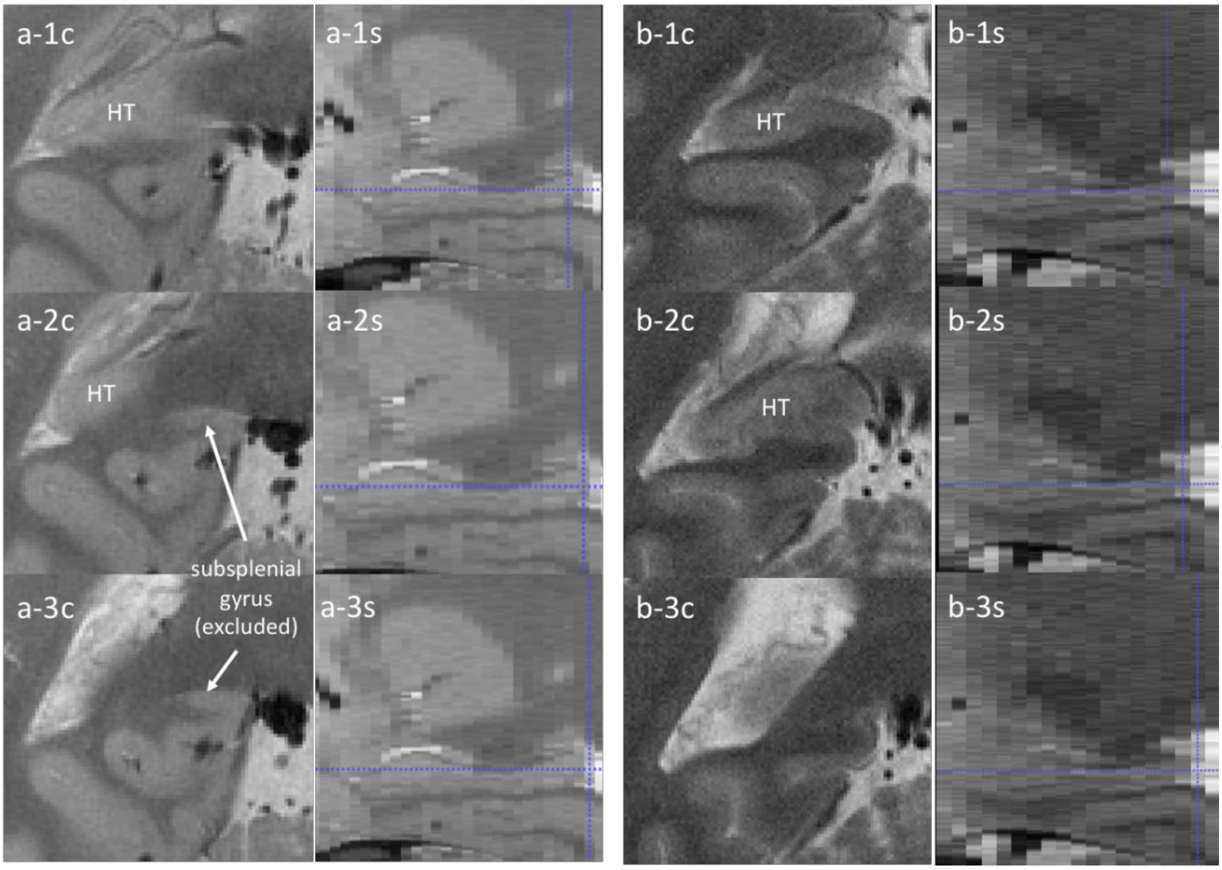

### 3. Ventral boundary

The HT is bordered ventrally by the white matter of the fusiform and lingual gyri as well as the cingulum bundle (Oishi et al., 2012). Consistent with the outer boundaries protocol of the hippocampal body (see Appendix B of the supplemental doc/pdf), the ventral boundary of the HT is defined by the border between the gray matter of the hippocampus and the white matter located inferior to it, such that white matter is excluded from the definition of the HT (Figure 3).

**Figure 3: Ventral boundary**
Three HT slices - anterior (1) to posterior (3) - shown for two different subjects (a and b) with label ("l") and unlabeled ("u").

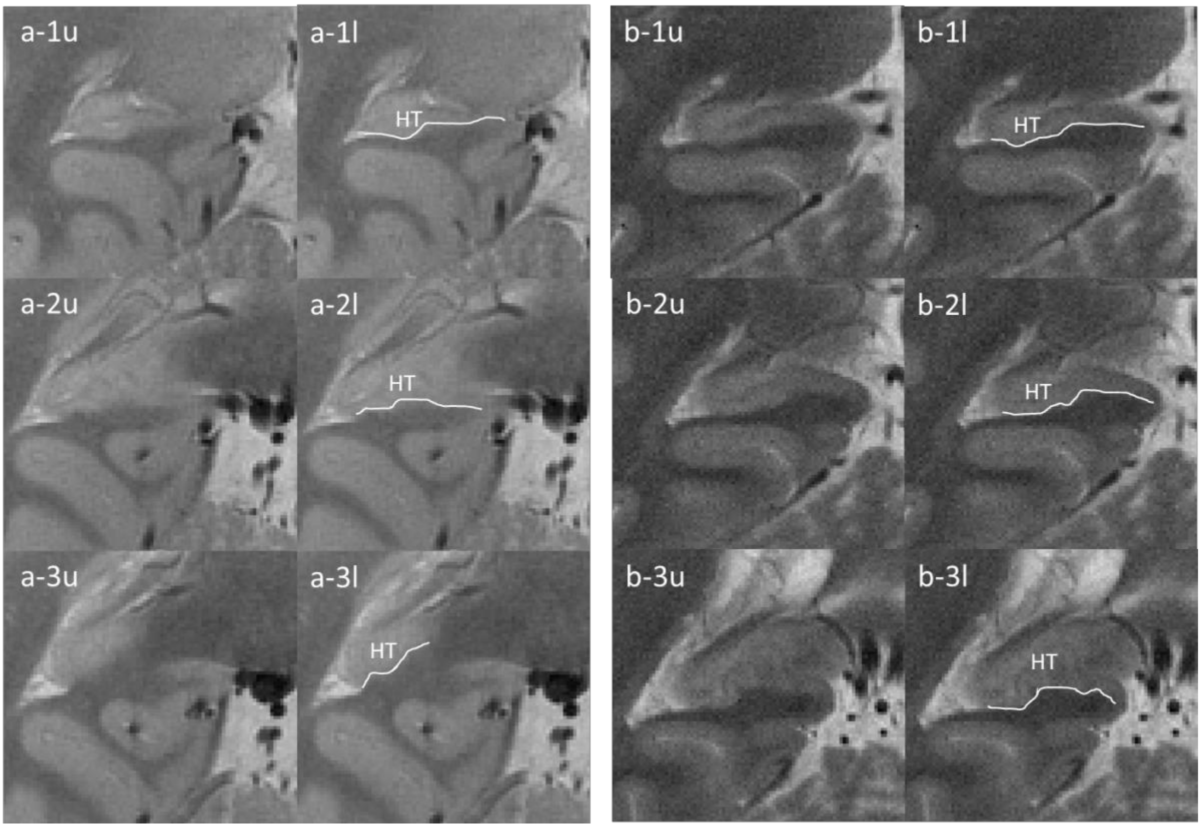

### 4. Dorsal boundary

The dorsal boundary is defined by the fimbria/fornix, the white matter of the splenium of the corpus callosum and/or CSF (see Figure 4). For some cases, a small piece of gray matter appears to detach from the main portion of the HT (see the dotted line in Figure 4 b-2). This detached part should not be considered HT as it could constitute either hippocampal or thalamic tissue. If the gray matter is not detached, then include the entire superior portion of the HT (as in Figure 4 a). Note that the fimbria and the fornix should not be included in the segmentation.

**Figure 4: Dorsal boundary**
Three HT slices - anterior (1) to posterior (3) - shown for two different subjects (a and b) with label ("l") and unlabeled ("u"). The dotted line demarcates a detached part of gray matter that should not be considered HT (see text). SCC: splenium of the corpus callosum

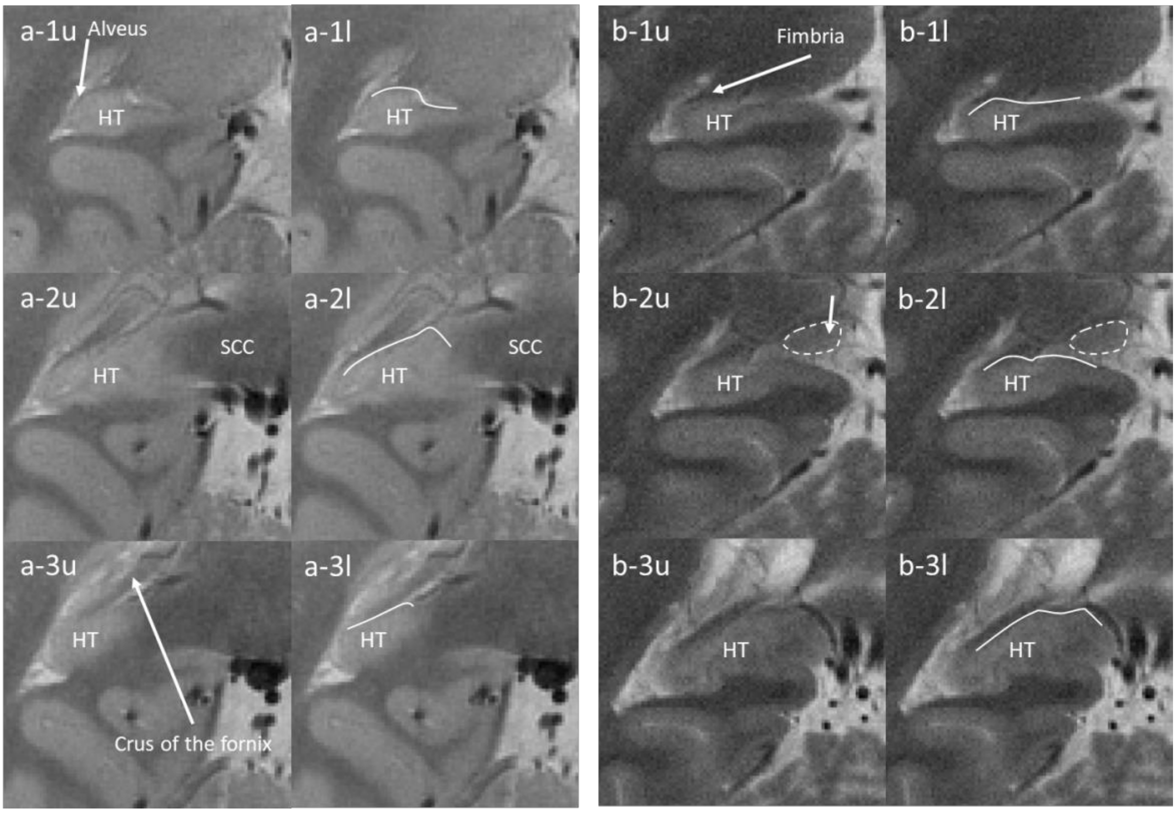

### 5. Lateral boundary

The lateral boundary of the HT is defined by the alveus and/or CSF if the alveus is not visible, such that the alveus should not be included in the segmentation (Figure 5). Partial voluming may be observed in this region given the mixture of signal stemming from a variety of proximal tissues/structures (gray matter, white matter, CSF, and choroid plexus), and so should be considered as part of the HT. Consistent with the recommendations in the outer boundaries protocol, contiguous slices should be consulted to evaluate whether the portion of tissue falls within the hippocampal region.

**Figure 5: Lateral boundary**
Three HT slices - anterior (1) to posterior (3) - shown for two different subjects (a and b) with label ("l") and unlabeled ("u").

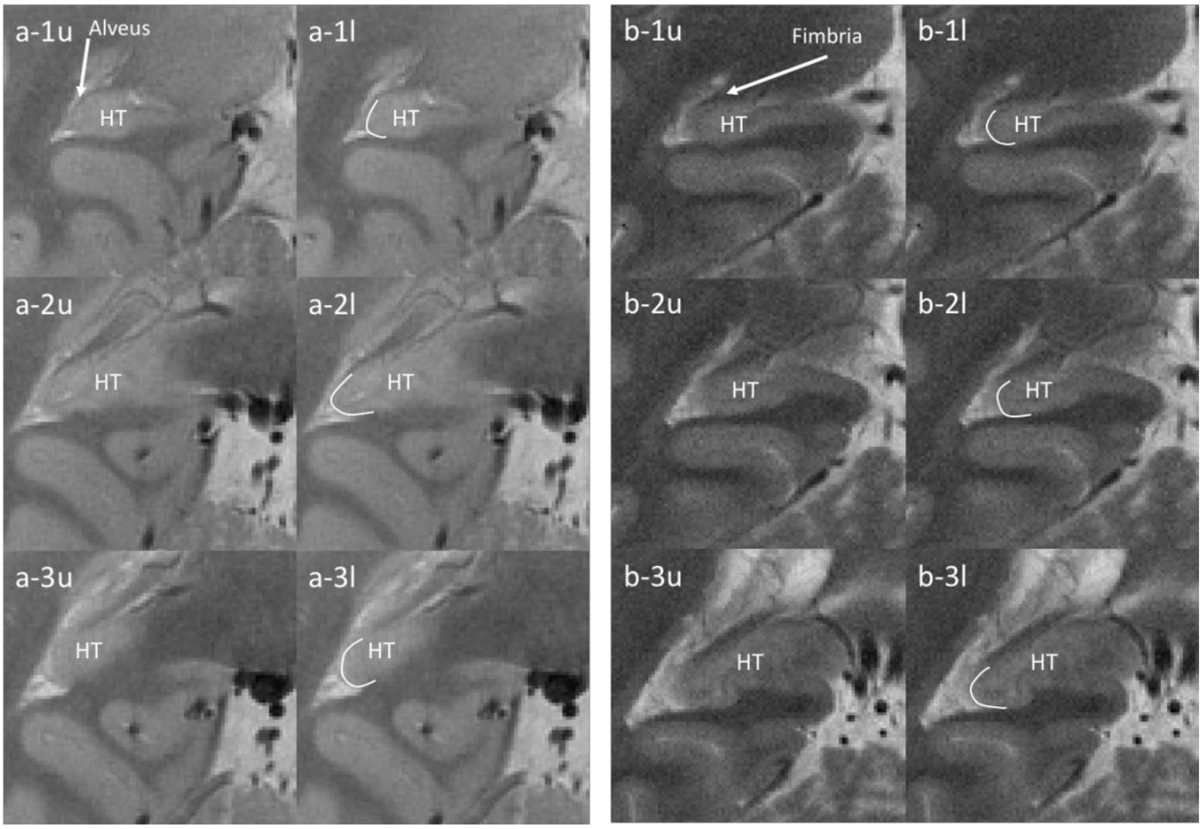

### 6. Medial boundary

Consistent with the outer boundaries protocol of the hippocampal body (see Appendix B of the supplemental doc/pdf), the medial boundary between the HT and parahippocampal cortex is defined as the most supero-medial corner of the parahippocampal gyrus (See the red dots in Figure 6 a-1, a-2, b-1, b-2, b-3). The boundary is placed at the level of maximum curvature in the cortical ribbon.

The HT blends with a gyrus sometimes referred to as subsplenial gyrus (Duvernoy et al., 2013). Given that the subsplenial gyrus contains a mixture of CA1 and CA3, this gyrus is segmented together with the HT and considered the same structure, as long as these two regions are connected. When the HT detaches from the subsplenial gyrus (Mai et al., 2008) (Figure 2 a-2c, a-3c ; Figure 6 a-3), the medial portion is excluded and only the lateral (mostly oval shaped) part of the HT is segmented. More medially or when the HT is detached from the subsplenial gyrus (Figure 6 a-3), the boundary is defined by the border between the gray matter of the hippocampus and the white matter.

**Figure 6: Medial boundary**
Three HT slices - anterior (1) to posterior (3) - shown for two different subjects (a and b) with label ("l") and unlabeled ("u"). For subject a, the HT detaches from the subsplenial gyrus (a-3u). SCC: splenium of the corpus callosum

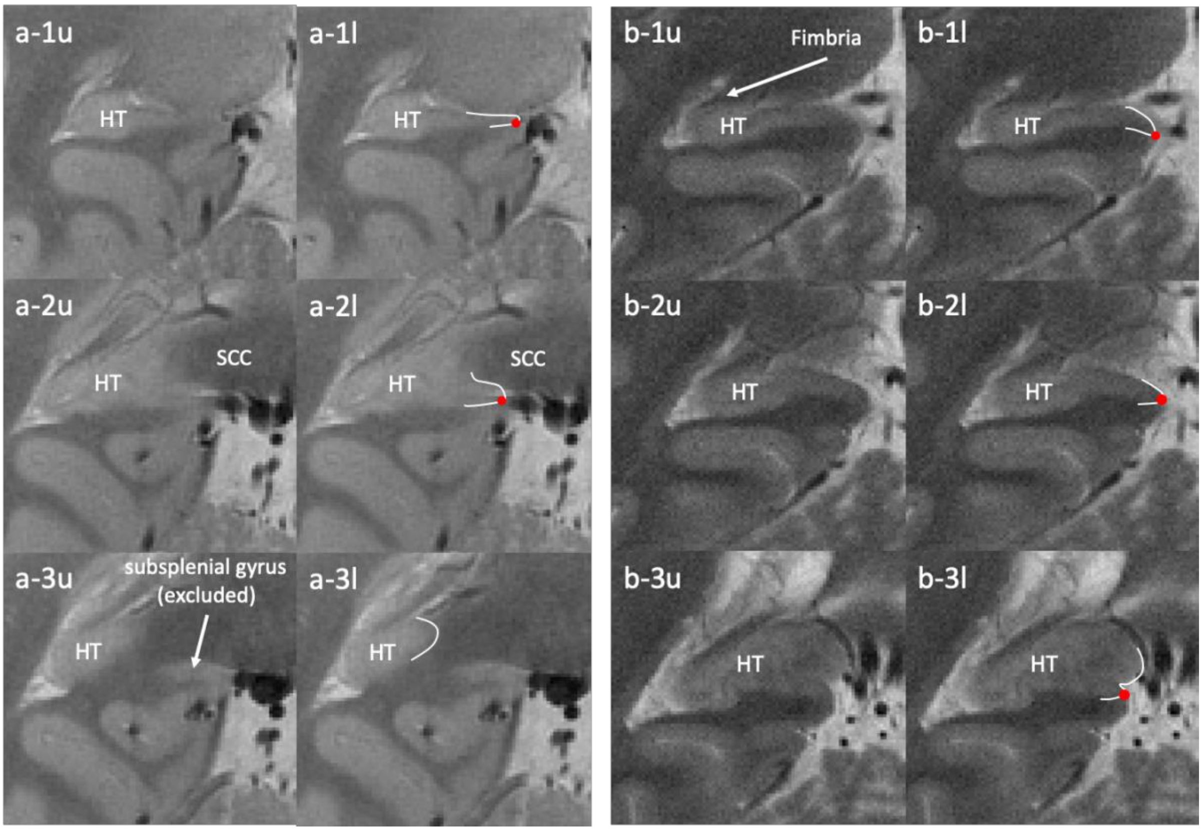

### 7. References

Duvernoy, H.M., Cattin, F., Risold, P.-Y., Vannson, J.L., Gaudron, M., 2013. The Human Hippocampus: Functional Anatomy, Vascularization, and Serial Sections with MRI. Springer

Mai, J. K., Voss, T., & Paxinos, G. (2008). *Atlas of the human brain*. Amsterdam: Elsevier/Academic Press.

Oishi, K., Faria, A. V., van Zijl, P. C. M., & Susumu Mori. (2012). *MRI Atlas of Human White Matter* (2nd ed.). Academic Press.

**Appendix B: Description of Outer Boundaries**

**Outer boundaries protocol**

Ana Daugherty and Renaud La Joie

On behalf of the Boundary Working Group

Last updated: July 2019

---

**Table of contents**

I. Anterior
II. Posterior
III. Dorsal
IV. Ventral
V. Medial
VI. Lateral
VII. Blood vessels
VIII. CSF and cysts

---

**Notes:**

All protocol rules relate to viewing the hippocampal body in the coronal view, and assume that boundaries will be drawn independently for the left and right hemispheres (such that asymmetries in boundary placement are possible).

All figures are T2 coronal images acquired perpendicular to the longitudinal axis of the hippocampus and are centered on the hippocampal body, unless otherwise specified.

#### I. ANTERIOR

The disappearance of the uncus is used to determine the transition from hippocampal head to body. The uncus lies on the medial edge of the hippocampal head, and on posterior head slices, it is often connected to the rest of the hippocampus via the fimbria only (See FIG 1). The anterior boundary of the hippocampal body (HB), i.e., the anterior-most slice included in the HB, is the **first slice posterior to the last visualization of the uncus**.

Note that head misalignment and/or anatomical differences between hemispheres may lead to differences in defining the anterior boundary of the HB in each hemisphere (usually differing by one slice in images with 2mm slice thickness). See FIG 1 and FIG 2 for examples.

Importantly, conservative judgement should be used and slices showing partial voluming of the uncus should be categorized as a “head slice” and not part of the HB. Partial voluming refers to situations in which a voxel represents an average of two or more tissue types (e.g., both grey matter and cerebrospinal fluid). When determining partial voluming, consult the previous contiguous slice to evaluate if the portion of tissue falls within the uncus region. See FIG 3 for an example.

**FIG 1**. Top left: Hippocampal head displaying the uncus in both hemispheres. Top right: Slice showing the disappearance of the uncus in the left hemishphere, and as such, the anterior-most slice of the hippocampal body in the left hemisphere. T2-weighted image, resolution 0.39 x 0.39 x 2mm. Bottom: Sketch adapted from Duvernoy et al., 2013 with portions of the uncus labeled as 4 and 6

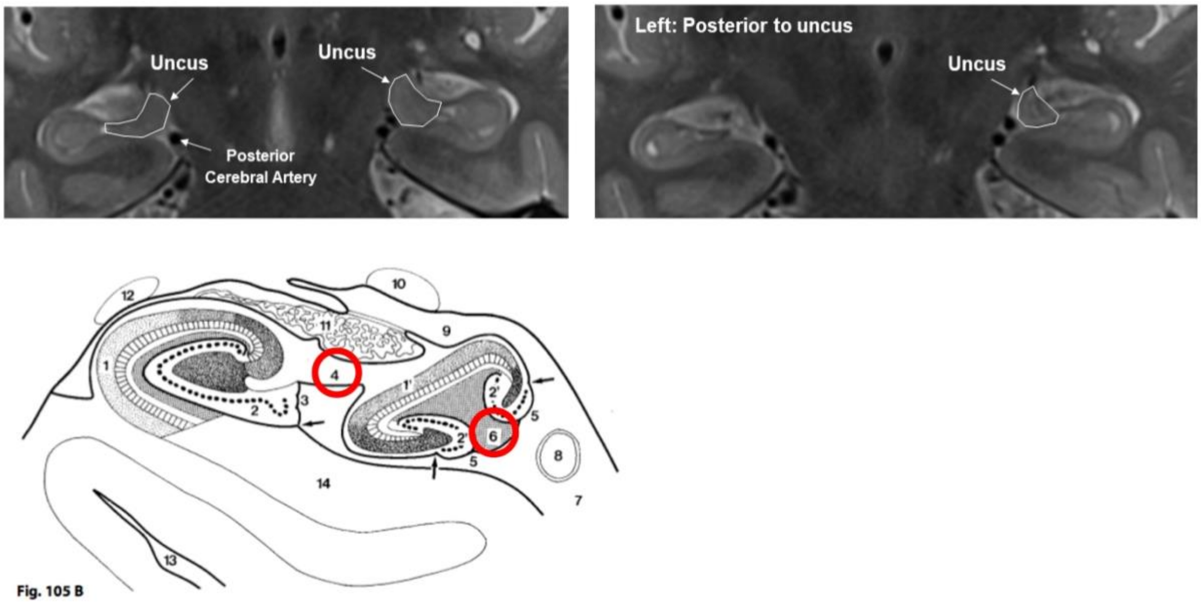

**FIG 2**. Second example demonstrating hemispheric asymmetry in the definition of the anterior boundary of the hippocampal body. Slices proceed from anterior to posterior. T2-weighted image, resolution 0.39 x 0.39 x 2mm.

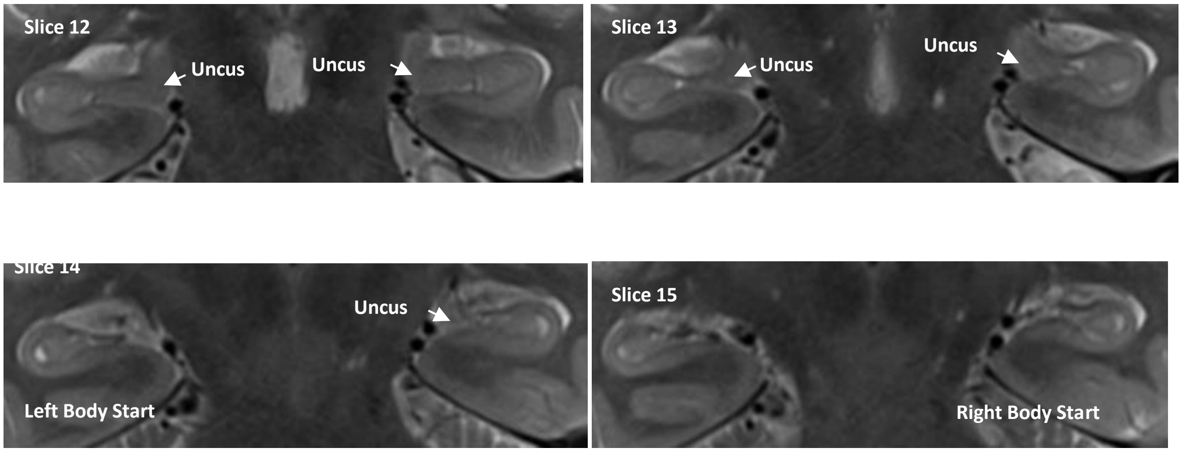


**FIG 3**. Example demonstrating partial voluming of the uncus. Slices proceed from anterior to posterior. T2-weighted image, resolution 0.39 x 0.39 x 2mm.

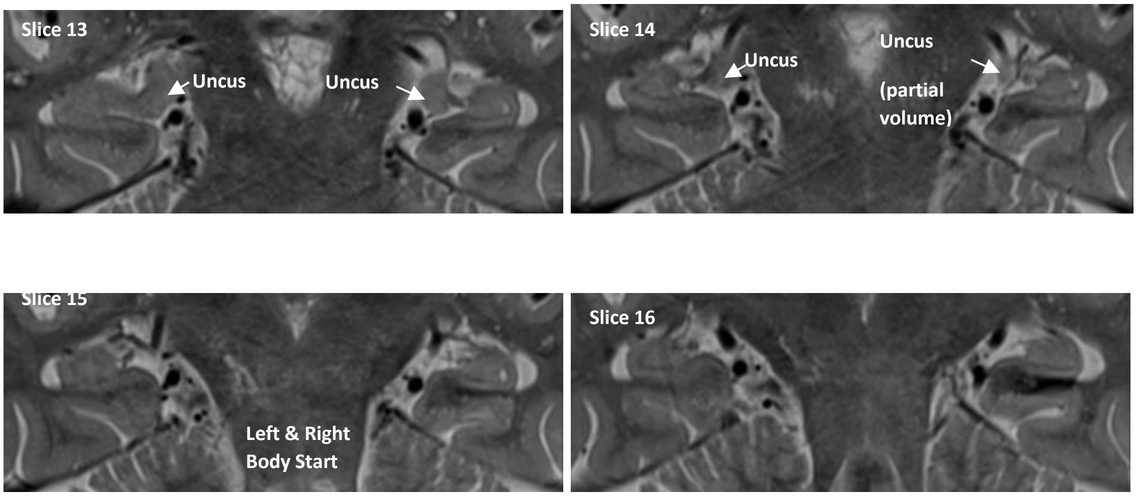

**FIG 4**. Example demonstrating partial voluming of the uncus (slice 10) on an image with motion artifacts. Slices proceed from anterior to posterior. T2-weighted image, resolution 0.39 x 0.39 x 3mm.

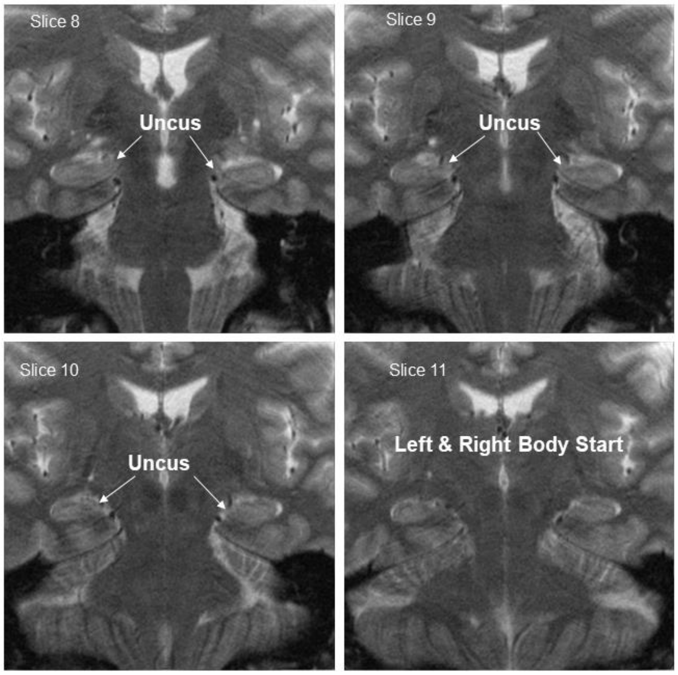


⇒ Back to table of contents

#### II. POSTERIOR

The posterior boundary of the HB, i.e., the posterior-most slice included in the HB, is determined by a single landmark: **the posterior-most slice on which the colliculi are visible**. The colliculi, also called the lamina quadrigemina, refer to the superior and inferior colliculi of the midbrain. On a T2-weighted coronal image, they appear as four hypointense (i.e., dark) round structures along the midline of the brain, posterior to the cerebral peduncles. When all four colliculi are visualized in the coronal plane, their arrangement may appear like a "butterfly". On 2-mm coronal slices, the colliculi will be visualized on 2-3 slices (see FIG 5 and FIG 6 for examples).

Segmentation of the subfields within the hippocampal body only stops when the colliculi have entirely disappeared. Note that partial voluming of the colliculi may occur, as shown in FIG 7-9, and that any visualization -- even partial -- should be considered a body slice when defining the posterior boundary of the HB.

The image acquisition of the hemispheres may be misaligned, and similar to the anterior-most slice, the posterior-most slice of the hippocampal body may differ between hemispheres. Only one colliculi in a hemisphere need be visualized to be considered the most-posterior slice of body in that hemisphere. It is worth noting that because the colliculi are structures that fall along the midline, this landmark will be less sensitive to differences between hemispheres as the anterior ranging rule. The use of this landmark is a conservative posterior definition of the HB, ensuring exclusion of the tail. In some cases, this may result in a portion of the HB mislabeled as hippocampal tail.

**FIG 5**. Left: Final posterior slice of the hippocampal body displaying the colliculi. This landmark coincides with others that are commonly reported in the literature: the crus fornix and the "tear drop" shape of the hippocampal body. Right: Colliculi are no longer visible, and as such this slice is considered the first slice of the hippocampal tail. T2-weighted image, resolution 0.39 x 0.39 x 2mm.

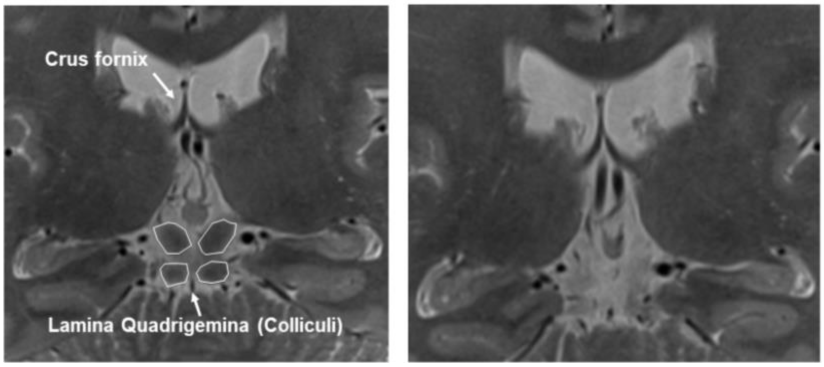

**FIG 6**: Series of coronal slices proceeding from anterior to posterior, demonstrating the definition of the posterior boundary of the hippocampal body. T2-weighted image, resolution 0.39 x 0.39 x 2mm.

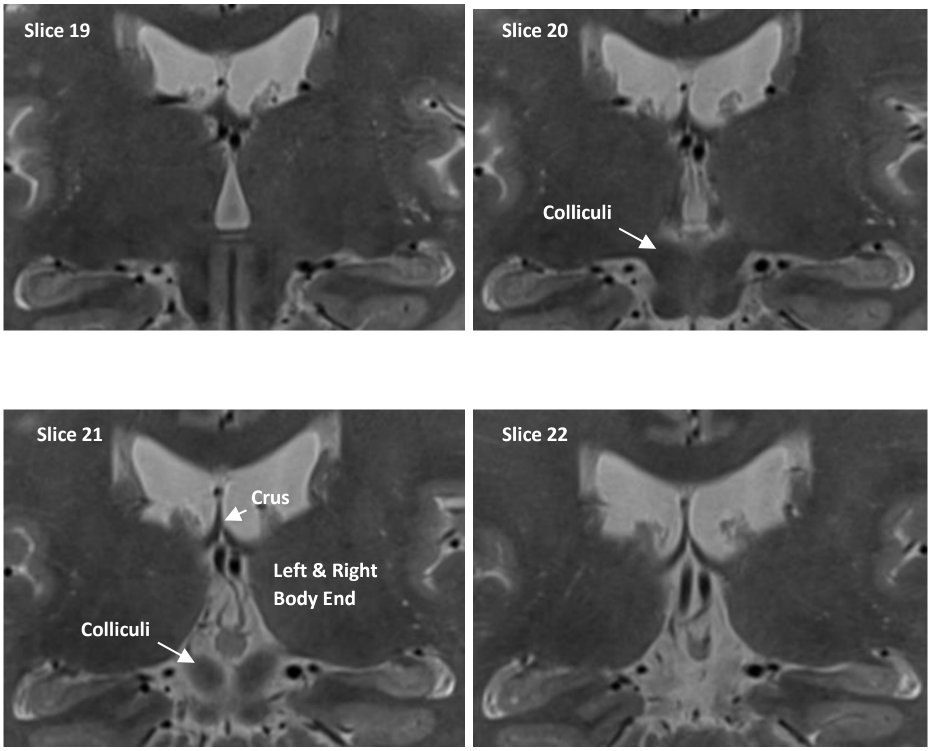


**FIG 7**: Series of coronal slices proceeding from anterior to posterior, demonstrating partial voluming of the colliculi and how even a partial visualization of the colliculi should be interpreted as a body slice. Note that the superior colliculi in each hemisphere are visualized with a portion of dark tissue that is consistent with the prior slice and this is interpreted as partial voluming. As such, the most-posterior body is on slice 22. T2-weighted image, resolution 0.39 x 0.39 x 2mm.

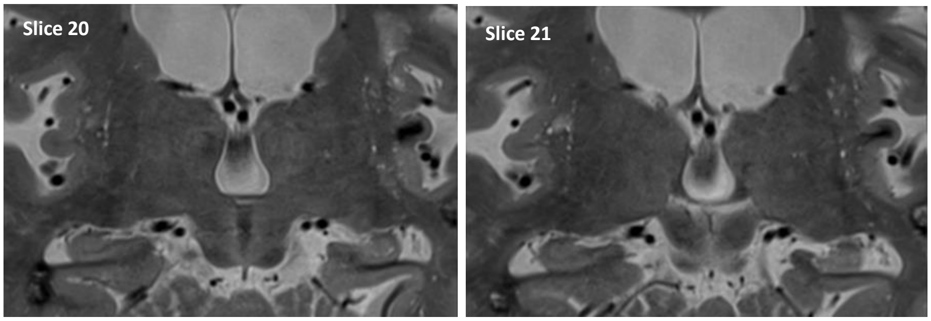

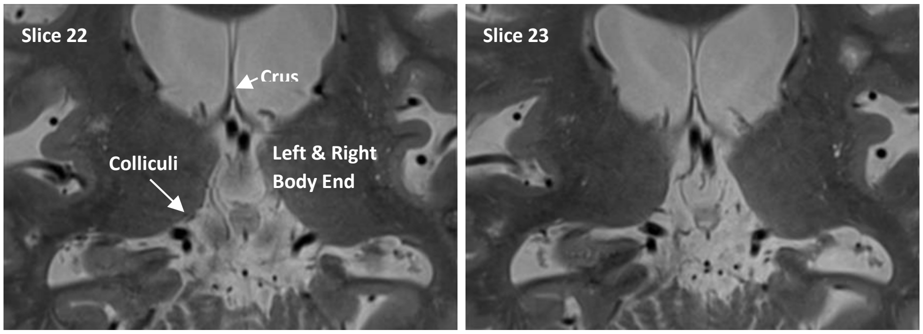


**FIG 8**: Series of coronal slices proceeding from anterior to posterior, demonstrating partial voluming of the colliculi with motion arifact. Note that the inferior colliculi in each hemisphere are visualized with a portion of dark tissue that is consistent with the prior slice and this is interpreted as partial voluming. As such, the most-posterior body is on slice 15. T2-weighted image, resolution 0.39 x 0.39 x 3mm.

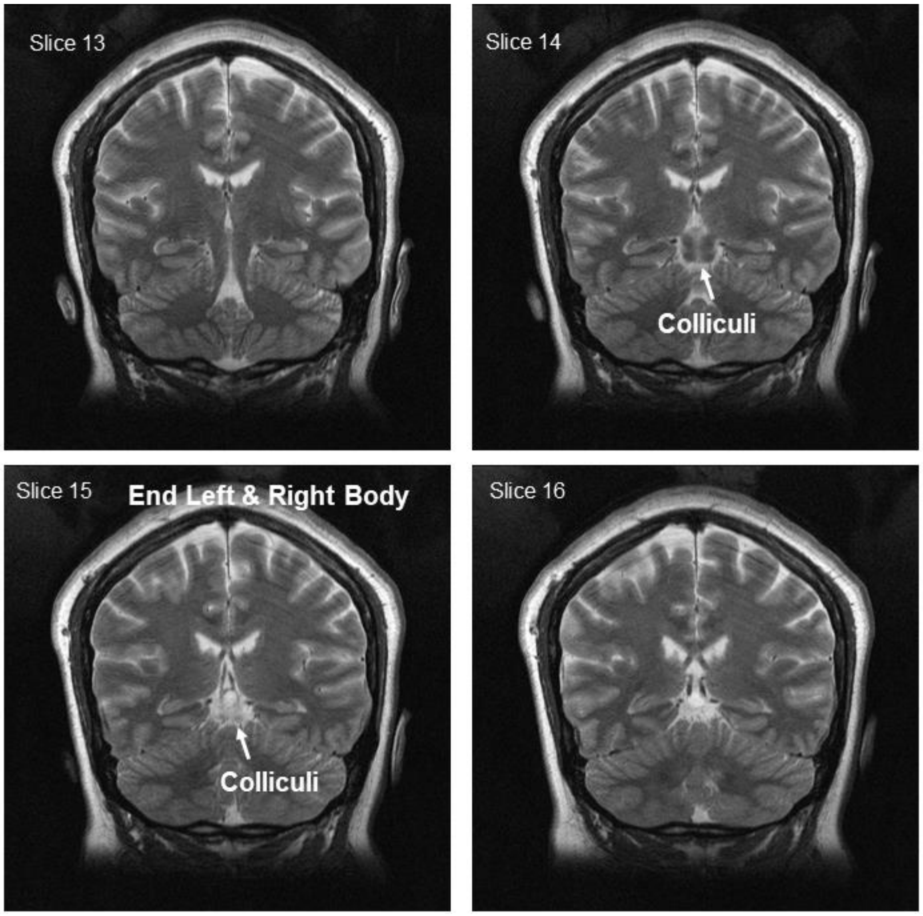

**FIG 9**: Series of coronal slices proceeding from anterior to posterior, demonstrating partial voluming of the colliculi with head misalignment at acquisition. Note that the inferior colliculi in each hemisphere are visualized with a portion of dark tissue that is consistent with the prior slice and this is interpreted as partial voluming. As such, the most-posterior body is on slice 23. T2-weighted image, resolution 0.39 x 0.39 x 2mm.

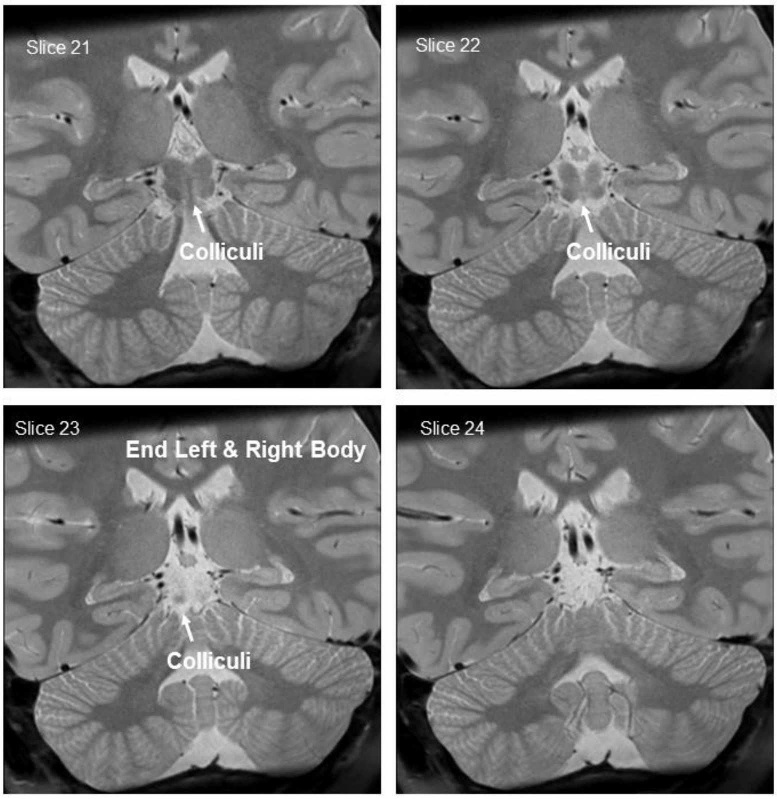


The boundary definition should be made foremost by visibility of the colliculi, even in instances of hemispheric or subtle head pitch misalignment. As can be seen in FIG 5-7, the posterior-most slice on which the colliculi are visualized often (but not always) shows the crus fornix and the HB often has the typical "tear drop" shape on this slice. If HB-like slices--on which the dentate gyrus is still visualized and the hippocampus has a tear drop shape--appear posterior to the last visualization of the colliculi, the researcher may suspect scan misalignment. On a scan aligned to be perpendicular to the hippocampal long axis, the inferior and superior colliculi will be visualized at the same time within the slice sequence, whereas on a misaligned scan, the inferior or superior colliculi (depending upon the direction of pitch) will be visualized out of sync. If the image acquisition is misaligned, and the image cannot be resliced to correct the alignment, then additional landmarks may be considered when defining the posterior boundary of the HB. The appearance of the crus of the fornix concurrent with a “tear drop” like shape of the hippocampus (including visualization of the dentate gyrus) should be considered the posterior-most slice of the hippocampal body.

⇒ Back to table of contents

##### III. DORSAL

The dorsal boundary is defined as the interface between the gray matter tissue of the HB and the white matter of the alveus and fimbria. In contrast to the low-intensity band of white matter sitting on top of the HB, the gray matter appears as high-intensity voxels on T2 MRI.

FIG 10: The alveus and fimbria in a coronal view in the anterior hippocampal body. Left: Sketch adapted from Duvernoy, 2005. Right: T2 MRI, image resolution: 0.42 x 0.42 x 2mm.

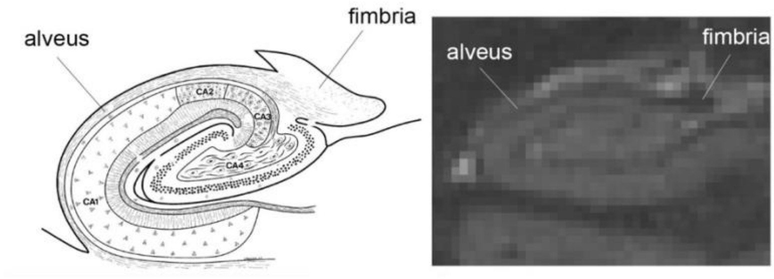


With respect to subfield segmentation, **both the alveus and fimbria must be excluded from any regions of interest**. They receive inputs from different hippocampal subregions and cannot be clearly assigned to one specific subregion. In addition, their inclusion would pose additional segmentation issues at the level of the crux of the fornix that would also contribute to reliability issues.

Therefore, the **hypointense (dark) voxels belonging to the fimbria** as well as the **thin hypointense band covering the hippocampal body on the dorso-lateral boundary** corresponding to the alveus **have to be excluded.**

FIG 11: Coronal view of the hippocampal body demonstrating the dorsal boundary along the anterior-posterior axis. The red line indicates the dorsal-most voxels to be included in the HB. Note: In the posterior part of the hippocampal body, the dentate gyrus extends medially. This extension has to be included in the hippocampal segmentation (panels D and E). Image resolution: 0.42 x 0.42 x 2mm.

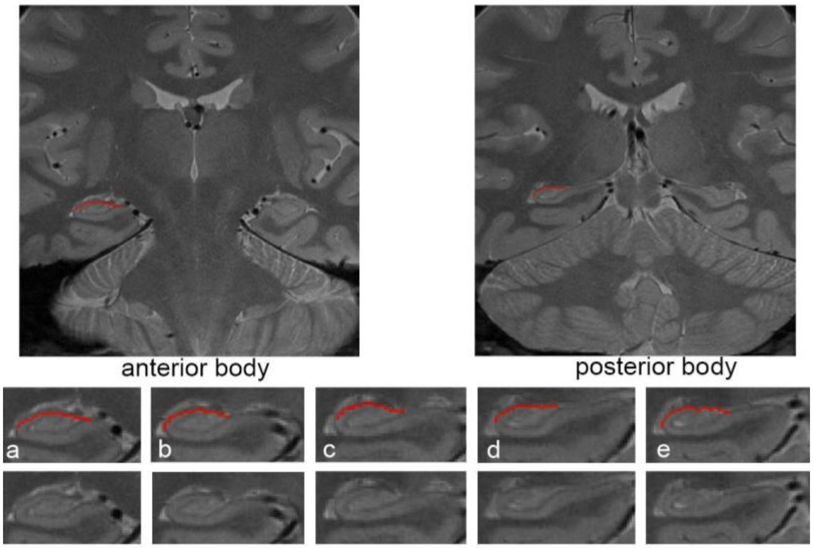


In contrast, the **hypointense voxels bordering the subiculum superiorly, and continuous with the stratum-lacunosum moleculare (SRLM) - are considered part of the hippocampus and included in the segmentation.**

**FIG 12**: Coronal view of the hippocampal body depicting incorrect and correct labeling of the hypointense voxels bordering the subiculum. Image resolution: 0.42 x 0.42 x 2mm.

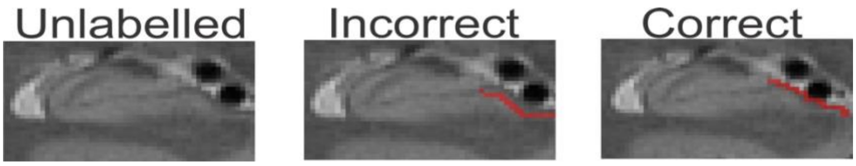


⇒ Back to table of contents

#### IV. VENTRAL

At the level of the HB, the ventral hippocampus is bordered by the white matter sitting above the parahippocampal gyrus. Following this logic, the ventral boundary is simply taken to be the **border between the grey matter of the hippocampus and the white matter that occurs inferior to it**, such that white matter is excluded from the definition of the HB.

**FIG 13**: Ventral boundary depicted in a coronal view of the hippocampal body. The red line indicates the ventral-most voxels to be included in the definition of the HB. Image resolution: 0.42 x 0.42 x 2mm.

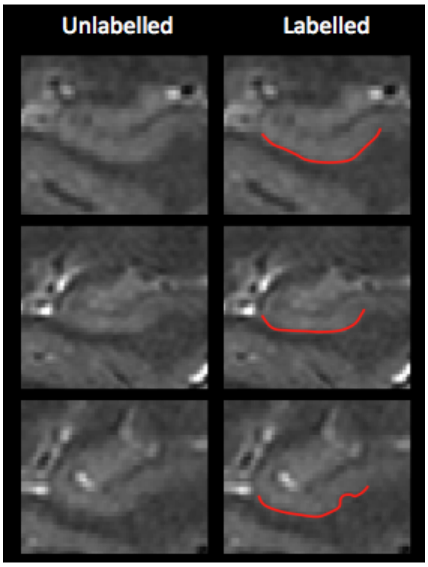


**FIG 14**: Coronal view of the hippocampal body demonstrating ventral boundary, adapted from Duvernoy et al., 2013.

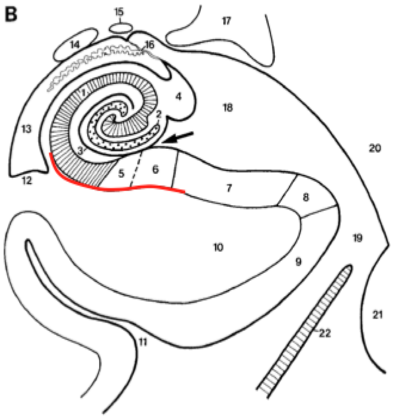


⇒ Back to table of contents

#### V. MEDIAL

At the level of the HB, the medial portion of the hippocampus corresponds to the subiculum and must be separated from the entorhinal, perirhinal and parahippocampal cortices. This separation is inherently difficult since few morphological or contrast differences exist to aid in differentiating the subiculum from the entorhinal/perirhinal/parahippocampal cortices.

The EADC-ADNI Harmonized Protocol (HaRP, http://www.hippocampal-protocol.net) defines the medial boundary by '*tracing an irregular line continuing from the visible interface with the WM of the parahippocampal gyrus, to the ventro-medial aspect based on the continuity of the boundary as detected from morphological details and GM intensity*'.

**FIG 15**: Top: **HaRP (<u>not HSG!</u>) medial boundary** depicted on a coronal MRI slice at the level of the hippocampal body. Bottom: HaRP medial boundary depicted on a coronal sketch of the hippocampal body adapted from Duvernoy et al, 2013. Image resolution: 0.42 x 0.42 x 2mm.

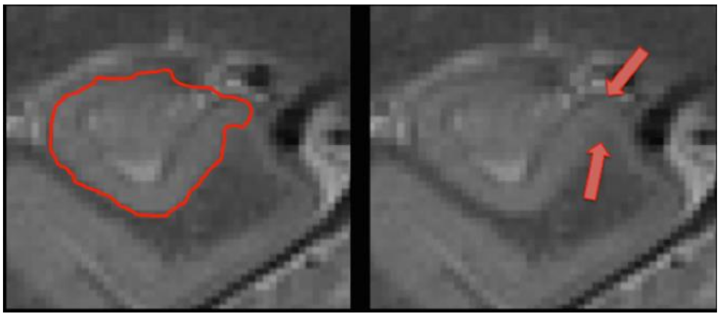

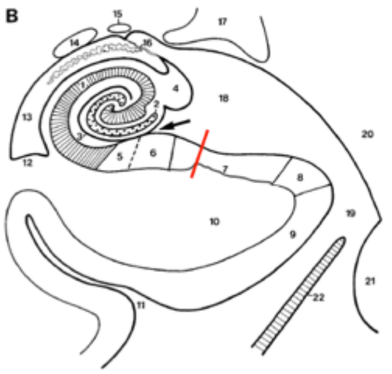


1. Cornu ammonis
2. Gyrus dentatus
3. Hippocampal sulcus (deep or vestigial part)
4. Fimbria
5. Prosubiculum
6. Subiculum proper
7. Presubiculum
8. Parasubiculum
9. Entorhinal area
10. Parahippocampal gyrus
11. Collateral sulcus
12. Collateral eminence
13. Temporal horn of the lateral ventricle
14. Tail of caudate nucleus
15. Stria terminalis
16. Choroid fissure and choroid plexuses
17. Lateral geniculate body
18. Lateral part of the transverse fissure
19. Ambient cistern
20. Mesencephalon
21. Pons
22. Tentorium cerebelli

The HSG protocol defines the medial boundary of the hippocampus differently than HaRP. Namely, at the level of the HB, **the medial boundary is defined as the most <u>medial</u> corner of the parahippocampal gyrus** and typically coincides with the termination of the subicular complex. See Figs 16-19 for examples. The boundary is placed at the **level of maximum curvature in the cortical ribbon**, just before it runs parallel to the tentorium cerebelli (i.e. the meninges indicated by blue arrows in Fig 16). Note that the boundary is traced perpendicular to the cortical ribbon (see Fig 19). It is important to note that this boundary **would more likely allow for the inclusion of presubicular and parasubicular cortices** than the HaRP protocol (see lower panel of Fig 16).

Depending on the orientation of the image acquisition and/or the shape of the hippocampus, the location of this boundary may be ambiguous. If, for example, the hippocampus has a rounder shape as in the top portion of Fig 16, and shows multiple areas of curvature, the medial boundary of the hippocampus is set at the medial point, where the cortical ribbon curves downwards and sometimes laterally, just before running parallel to the tentorium cerebelli.

FIG 16: Top: **Current protocol's definition of the medial boundary** at the level of the hippocampal body on a coronal MRI slice. The medial boundary (red arrow) is defined as the most medial point of the cortical ribbon before the cortex runs parallel to the tentorium cerebelli (meninges, indicated by the blue arrows). Bottom: Current protocol's definition of the medial boundary displayed on a coronal sketch of the hippocampal body from Duvernoy et al, 2013. Image resolution: 0.42 x 0.42 x 2mm.

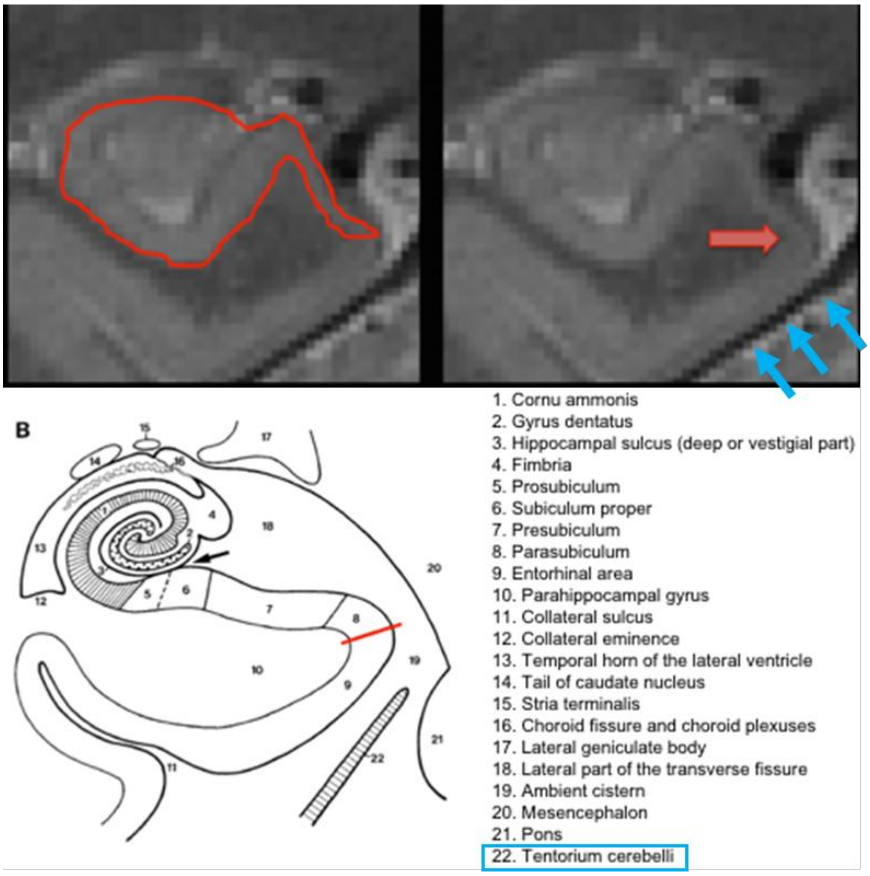


At times, the observed thinning of the subicular cortex and lower MRI contrast may limit tracing ability, yet most high resolution images offer the spatial resolution to complete tracing to this most medial point.

Depending on the subject anatomy and the image orientation, the most medial aspect of the medial temporal lobe can be flat (~vertical, see Subjects 2 and 3 from figure 17l) and the level of maximal curvature might correspond to the most supero-medial point of the parahippocampal gyrus.

**FIG 17**: Examples of defining the medial boundary at the level of the hippocampal body. The medial boundary is defined as the area of maixma curvature of the parahippocampal gyrus (most medial point for Subject 1; or superomedial corner in case the most medial aspect of the medial temporal lobe is flat/vertical, see Subjects 2-3), where the cortex curves downwards before running parallel to the tentorium cerebelli. Coronal slices progress from anterior (top row) to posterior (bottom row). Image resolution: 0.42 x 0.42 x 2mm.

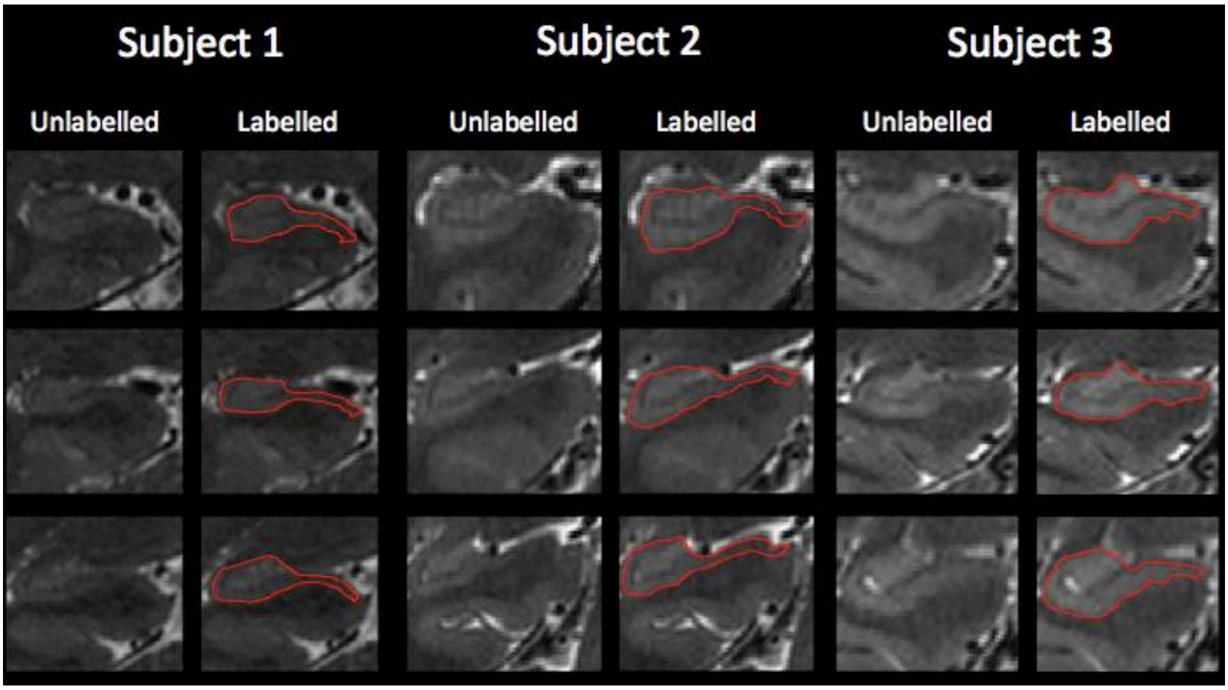


**FIG 18**: Examples of correct and incorrect labeling of the medial boundary depicted at the level of the hippocampal body. Correct labeling demonstrates that the boundary is placed at the most (supero)medial point of the parahippocampal gyrus, at its maximum curvature before turning downwards to run parallel to the tentorium cerebelli. Image resolution: 0.42 x 0.42 x 2mm.

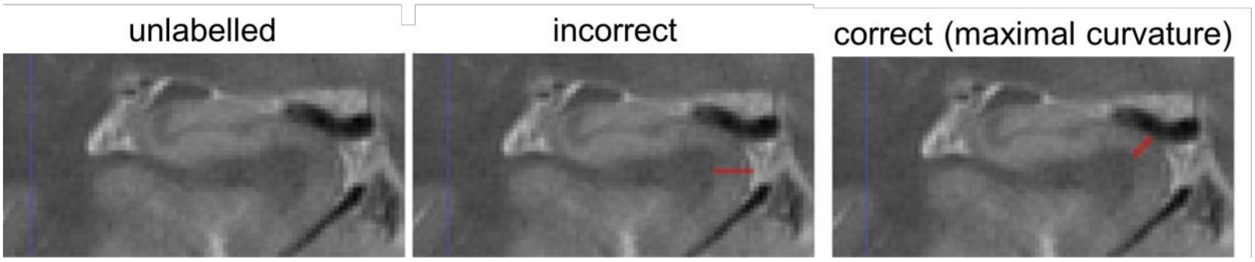


Furthermore, care should be taken to draw this border **orthogonal (perpendicular) to the cortical ribbon/pial surface**.

**FIG 19**: Segmentation of the medial boundary with cortical ribbon

highlighted to demonstrate correct orthogonal placement (the medial boundary of the hippocampus is set perpendicular to the dotted line that represents the pial surface at the level of maximal curvature)

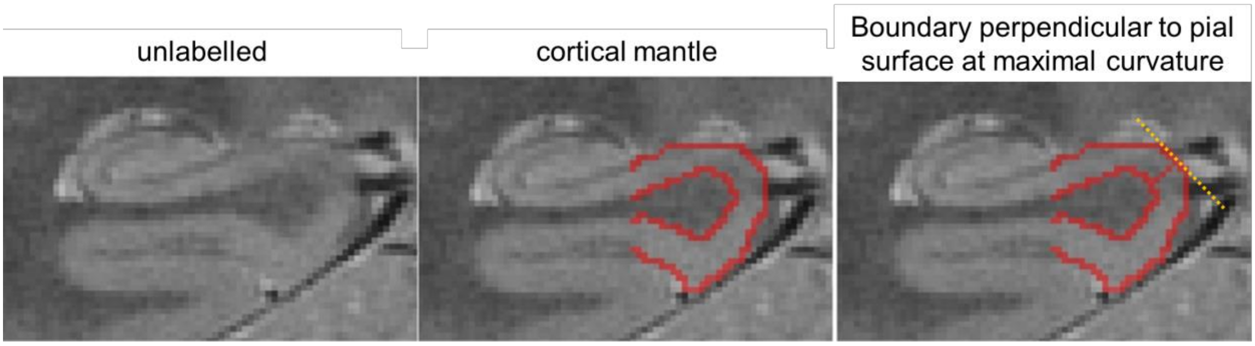


⇒ Back to table of contents

#### VI. LATERAL

The lateral boundary is the same as that presented in the "Dorsal Boundary" section. That is, the lateral boundary is taken to be the border between the gray matter tissue of the HB and the white matter of the alveus/fimbria, such that the **alveus and fimbria should not be included in the segmentation**.

**FIG 20**: Top: Examples of the lateral boundary drawn on several coronal slices at the level of the hippocampal body on MRI. Similar to the dorsal boundary, the lateral boundary is defined as the interface between the hippocampal grey matter and extra-hippocampal white matter. Image resolution: 0.42 x 0.42 x 2mm. Bottom: Lateral boundary displayed on a coronal sketch of the hippocampal body adapted from Duvernoy et al, 2013.

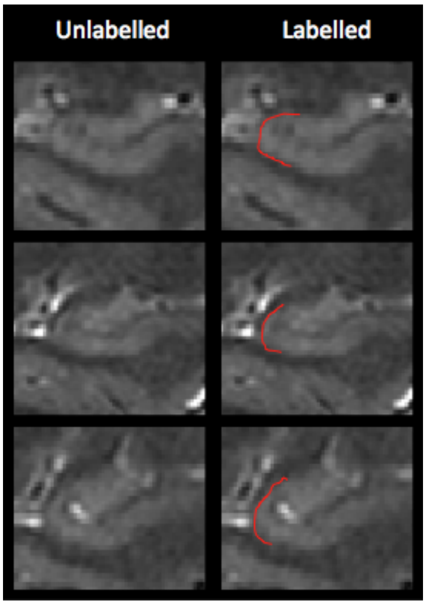


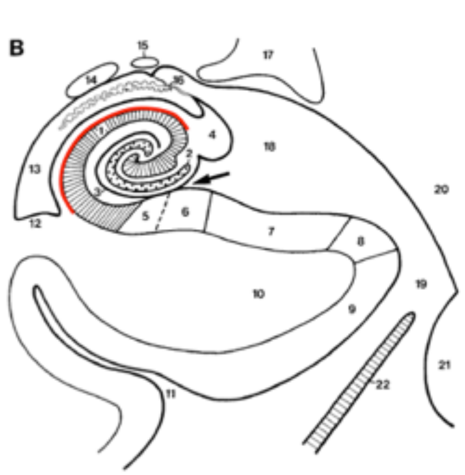


1. Cornu ammonis
2. Gyrus dentatus
3. Hippocampal sulcus (deep or vestigial part)
4. Fimbria
5. Prosubiculum
6. Subiculum proper
7. Presubiculum
8. Parasubiculum
9. Entorhinal area
10. Parahippocampal gyrus
11. Collateral sulcus
12. Collateral eminence
13. Temporal horn of the lateral ventricle
14. Tail of caudate nucleus
15. Stria terminalis
16. Choroid fissure and choroid plexuses
17. Lateral geniculate body
18. Lateral part of the transverse fissure
19. Ambient cistern
20. Mesencephalon
21. Pons
22. Tentorium cerebelli

⇒ Back to table of contents

#### VII. BLOOD VESSELS

There are several blood vessels within and close to the hippocampal formation. Of interest are especially the posterior cerebral artery and the basal vein.

**FIG 21**: Coronal sketch at the level of the hippocampal body depicting blood vessels neighboring the hippocampus. Adapted from Duvernoy, 2005.

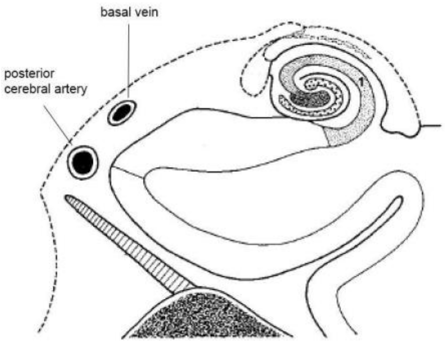


There are also smaller intrahippocampal branches from these vessels connecting the inner hippocampus. Blood vessels appear hypointense (dark) in T2 weighted images (note, however, that they are bright in T1 images). The big vessels close to the subiculum can be especially problematic during segmentation as they can cause signal drop out or may cause hippocampal anatomy to appear slightly different. **Vessels and/or signal drop out due to vessels has to be excluded from the segmentation.**

**FIG 22**: Two examples of arteries close or within the hippocampal body. The left column is unlabeled and the right column indicates proper segmentation according to the current protocol. In the first example (A and B), the artery is touching the grey matter, but in the second example (C and D) the artery appears within the grey matter and causes signal dropout. Image resolution: 0.42 x 0.42 x 2mm.

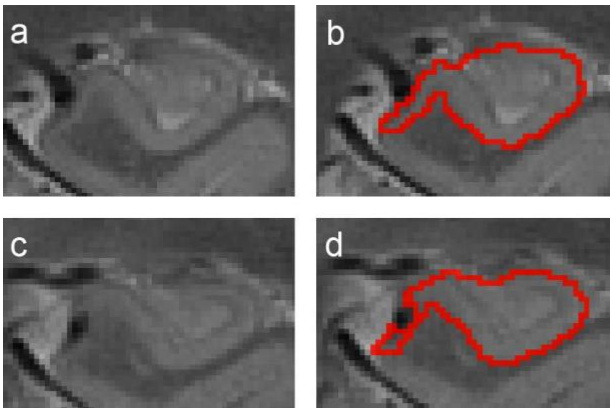


⇒ Back to table of contents

#### VIII. CSF and CYSTS

In T2 weighted images, CSF cysts appear as hyperintense (bright) regions (though note that they appear dark in T1 images). Cysts are observed regularly on higher resolution scans. They are often located in the vestigial sulcus along the longitudinal axis of the hippocampus, and a majority of them appear in the ventrolateral flexion points of CA1 (see below; also see Veluw et al., 2013).

In general **we recommend segmenting the cysts separately** either before or following the delineation of the other subfields. Cysts will be segmented using a separate label.

They **can often be followed on consecutive slices** [rule 1]. However, especially on images with anisotropic voxel size and thicker slices this does not have to be the case.

Cysts occurring in the HB **should be removed if the tracer is sure that they represent CSF**. Only clusters that consist of **at least two or more contiguous voxels (in any direction) that are considerably brighter than their surroundings will be labeled** [rule 2]. Often the voxels in the center of the cyst are brightest and fade out towards the edges (see FIG 13 from prior section).

Therefore, **cysts will be segmented based on hyperintensity, presence on adjacent slices, and number of voxels**. However, not every cyst might have all three properties.
**FIG 23**: Unlabeled (left column), incorrectly labeled (middle column), and correctly labeled (right column) CSF cysts. Slices are from the same subject and move from anterior (top row) to posterior (bottom row) portions of the hippocampal body. Note that for correct labeling, only the center, brightest voxels of the cysts are labeled as CSF. Image resolution: 0.42 x 0.42 x 2mm.

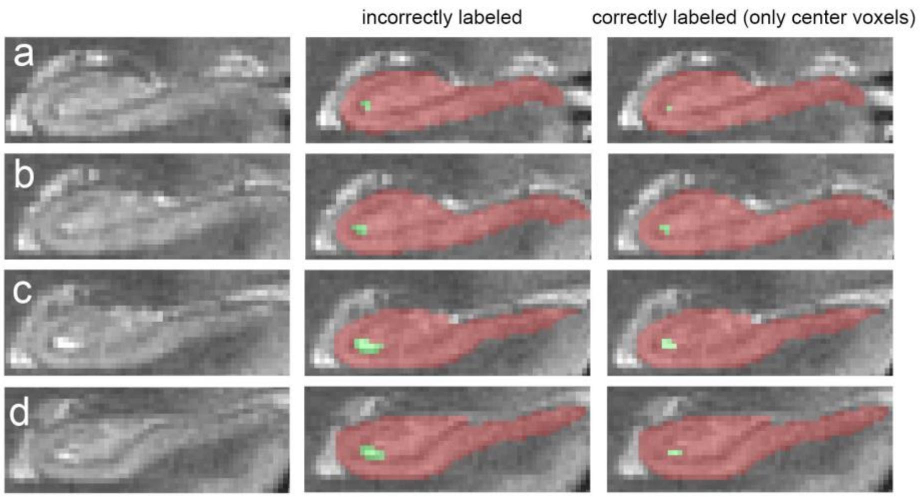


⇒ Back to table of contents
